# The dynamics of ACR and DNA methylation impact asymmetric subgenome dominance in allotriploid *Brassica* species

**DOI:** 10.1101/2025.02.16.638486

**Authors:** Chengtao Quan, Shengwei Dou, Cheng Dai

## Abstract

Polyploidy plays a crucial role in the evolution and diversification of eukaryotes, often leading to structural and functional imbalances as multiple subgenomes merge. Numerous studies have demonstrated the phenomenon of asymmetric subgenome dominance in polyploid species. However, the subgenomic dominance resulting from interspecific hybridization with accessible chromatin regions (ACRs) and DNA methylation remains largely unexplored in allotriploids. We generated two allotriploid hybrids of *Brassica* species (A_r_A_n_C_n_) by crossing the allotetraploid *Brassica napus* (A_n_A_n_C_n_C_n_) with the diploid *Brassica rapa* (A_r_A_r_). Our results revealed that the A_n_ subgenome exhibits the highest levels of gene expression among the three subgenomes (A_r_, A_n_, and C_n_) in F_1_ hybrids. However, the C_n_ subgenome that actually exerts dominance over A_n_ and A_r_, as evidenced by the abundance of dominant triplet homologous genes. We found that the C_n_ subgenome contains a significantly higher density of ACRs than A_n_ and A_r_ subgenomes, and the expression levels of dominant triplet homologous genes are closely associated with the presence of ACRs in proximal and genic. Interestingly, variations in DNA methylation levels alone do not fully explain the expression advantages observed in F_1_ hybrids. We then discovered that reduced CHH methylation in F_1_ hybrids might be associated with decreased expression of *BnaDCL3* and *BnaDRM2* in the RNA-directed DNA methylation pathway. Furthermore, the mutants *Bnadcl3^CR^* and *Bnardr2^CR^* exhibited a selective preference for 24-nt small RNA and non-CG methylation in the A_n_ and C_n_ subgenomes. Overall, this study elucidates the relationship between ACRs, DNA methylation, and asymmetric subgenome dominance in resynthesis allotriploid species.

## INTRODUCTION

Interspecific hybridization introduces multiple identical or different genomes into the nucleus for recombination, which results in either autopolyploidy or allopolyploidy (Doyle et al. 2008; Edger et al. 2018). In allopolyploids, it often possesses two or more structurally and functionally asymmetric subgenomes. The fusion of these multiple subgenomes can often lead to “genome shock,” which is an important factor in the process of evolution (Zhang et al. 2021b; Zhang et al. 2023; Li et al. 2023b). This phenomenon can induce diverse novel phenotypes in plants, increase their adaptability to various environmental conditions, and ultimately facilitate the emergence of new species (Cao et al., 2023b; Duan et al. 2023; He et al. 2022).

Interspecific hybridization often triggers rapid and extensive genomic reprogramming responses, significantly altering gene expression and epigenetic modifications (Jordan et al. 2020; Quan et al. 2022; Shao et al. 2019; Baldauf et al. 2022). Some of the immediate changes in gene expression following interspecific hybridization can be considered adaptive responses of the newly formed hybrid genome to the hybridization process. In polyploid plants, the dominant subgenome typically retains more genes, exhibits higher levels of gene expression, and experiences more intense purifying selection pressure-a phenomenon known as “genome dominance” (Alger et al. 2020; Bird et al. 2018; Glover et al. 2016). This phenomenon is commonly observed in polyploid plants, especially those formed by whole genome duplication (WGD) or hybridization (Zhang et al. 2021a; Zhang et al. 2021b; Wang et al. 2023). Dominant subgenomes have been well documented in species such as wheat, *Brassica napus*, and cotton (Ramírez-González et al. 2018; Chalhoub et al. 2014; You et al. 2023). In natural hexaploid wheat, the genome consists of three subgenomes (A, B, and D), with the D subgenome often displaying dominant traits (Yuan et al. 2020; Jordan et al. 2020). Genome-wide unbalanced expression has been noted in synthetic *Raphanus sativus-Brassica oleracea* intergeneric hybrids (Zhang et al. 2021a). However, the phenomenon of subgenomic dominance resulting from hybridization remains largely unexplored.

Epigenetics refers to various mechanisms that lead to long-lasting and inheritable changes in gene expression without modifying the DNA sequence (He et al. 2011). Key processes involved in epigenetics include chromatin remodeling, histone modifications, DNA methylation, and non-coding RNAs (Li et al. 2023a; Zhang et al. 2021b; Zhang et al. 2022). For instance, gene transcription is commonly regulated by the interactions between regulatory proteins and *cis*-regulatory elements (CREs) (Wang et al. 2020; Zhang et al. 2022; Liu et al. 2023), and the active CREs are found within active chromatin regions (ACRs) (Jordan et al. 2020). In *B. napus*, the C_n_ subgenome exhibits a higher level of chromatin accessibility compared to the A_n_ subgenome, which is influenced by the chromatin accessibility of specific genes within the C_n_ subgenome (Li et al. 2022). Similarly, the dominant subgenome shows increased levels of chromatin accessibility, while the loss of ACRs near certain genes is associated with decreased gene expression in maize (Yin et al. 2022). Recent research has revealed that DNA methylation in subgenomes changes dynamically during interspecific hybridization (Yuan et al. 2020). In *Juglans regia* and its wild relative *Juglans mandshurica*, gene pairs exhibit significantly elevated DNA methylation levels in the submissive subgenome relative to the dominant subgenome (Li et al. 2023b). The DNA methylation level in the promoter of the C_n_ subgenome is higher than that in the A_n_ subgenome in the synthetic allopolyploid *B. napus* (Bird et al. 2021). These results suggest that identifying ACRs and patterns of DNA methylation in plants has greatly improved our understanding of the intricate transcriptional regulatory networks that underpin genomic features and gene expression.

DNA methylation occurs through distinct pathways in CG, CHG, and asymmetric CHH (where H = A, T, or C) contexts (Law et al. 2010; He et al. 2011). In *Arabidopsis*, maintenance methylation of CG, CHG, and CHH is regulated by *METHYLTRANSFERASE1* (*MET1*) (Kankel et al. 2003), *CHROMOMETHYLASE3* (*CMT3*) (Lindroth et al. 2001), and *CHROMOMETHYLASE2* (*CMT2*) (Stroud et al. 2014), respectively. In an interspecific hybrid cross between *Arabidopsis thaliana* and *Arabidopsis lyrata*, a transposable element (TE)-specific increase in CHH methylation was observed at the chromosomal level in the hybrid compared to *A. thaliana* (Zhu et al. 2017). Two overlapping pathways regulate the CHH methylation of TEs, with *CMT2* predominantly functioning in heterochromatic regions and long transposons, accounting for approximately 70% of the total CHH methylation (Stroud et al. 2014). The remaining 30% of CHH methylation is maintained by *DOMAINS REARRANGED METHYLTRANSFERASE 2* (*DRM2*), which plays a pivotal role in the RNA-mediated DNA methylation (RdDM) pathway (Matzke et al. 2014). Recent findings indicate that subgenome DNA methylation undergoes dynamic changes during interspecific hybridization. In the case of *Juglans regia* and its wild relative *J. mandshurica*, gene pairs exhibit significantly elevated DNA methylation levels in the submissive subgenome relative to the dominant subgenome (Li et al. 2023b). The DNA methylation level in the promoter of the C_n_ subgenome was higher than that of the A_n_ subgenome in the synthetic *B. napus* (Bird et al. 2021). Mutations in the RdDM pathway have resulted in only subtle differences in DNA methylation in *Arabidopsis* (Yang et al. 2016). However, the impact of RdDM loss on subgenomic asymmetry in allopolyploid species remains to be elucidated.

The Brassicaceae family is vital for the global production of vegetable oil, animal feed, and vegetables for human consumption (Nikolov et al. 2019; Franzke et al. 2011). This family is an excellent resource for interspecific hybridization, which can create allopolyploids (Quan et al. 2022; Cao et al. 2023a; Cheng et al. 2016). Previous studies have investigated the gene expression and asymmetric dominance of different subgenomes in natural and resynthesized allopolyploid *Brassica napus* (A_n_A_n_C_n_C_n_). For instance, the expression levels of the majority of A_n_ subgenome genes show higher expression than C_n_ subgenome genes in *B. napus* pol CMS line 2063A, B409, and their F_1_ hybrid HZ62, the expression levels of most genes from the A_n_ subgenome are higher than those from the C_n_ subgenome across various tissues (Zhang et al. 2021b). In contrast, the expression levels of homologous gene pairs do not differ between A_n_ and C_n_ (Zhang et al. 2021b). However, in natural *B. napus* (*Darmor*) and resynthesized *B. napus* L. (HC-2 (maternal x paternal: *Brassica oleracea* 3YS013 x *Brassica rapa* 9JC002)), the homologous gene pairs in C_n_ subgenome are dominant over A_n_ in leaf tissue (Li et al. 2021). A similar pattern is also found in resynthesized *B. napus* (maternal x paternal: *Brassica oleracea* TO1000 x *Brassica rapa* IMB218) (Bird et al. 2021). These results indicate significant differences in the homologous expression levels of the A_n_ and C_n_ subgenomes of resynthesized *Brassica napus* derived from different parental lines. Further studies uncover that the differential imbalanced A_n_ and C_n_ subgenome expression of homologous genes is probably due to epigenetic modification (e.g., DNA methylation, chromatin accessibility, and histone modifications) (Li et al. 2021; Bird et al. 2021; Zhang et al. 2021b; Li et al. 2022).

Interspecific hybridization between *Brassica rapa* (A_r_A_r_) and *B. napus* (A_n_A_n_C_n_C_n_) is an effective way to expand the genetic base of *B. napus*. This approach leverages the significant potential of natural triploids in polyploid breeding (Leflon et al. 2010; Zhang et al. 2019). Moreover, these allotriploid hybrids offer a unique opportunity to investigate the effects of polyploidization on global gene expression, facilitating precise comparisons between F_1_ hybrids and their diploid or tetraploid progenitors. Previous studies on interspecific hybrids of *B. napus* and *B. rapa* have primarily concentrated on genomic structural variations and differences in gene expression (Cao et al. 2023b; Leflon et al. 2010; Qian et al. 2006; Zou et al. 2011; Zhang et al. 2019). However, the epigenetic regulatory differences arising from the interaction between the A_n_, C_n_, and A_r_ subgenomes remain unclear. In this study, we obtained a series of allotriploid hybrids (A_r_A_n_C_n_, 2n = 29) by crossing *B. rapa* (A_r_A_r_, 2n = 20) with two *B. napus* (A_n_A_n_C_n_C_n_, 2n = 38). We conducted a comparative analysis of DNA methylation, siRNA, accessible chromatin regions, and gene expression across the subgenomes of F_1_ hybrids, parentals, and *in silico* ‘hybrids’, revealing that the C_n_ subgenome exhibits dominance over the A subgenomes (A_n_ and A_r_) in resynthesized allotriploid F_1_ hybrids by comparing the number of dominant triplet homologous genes. Further, we noticed that the expression patterns of dominant and suppressed homologous genes were closely linked to the presence of ACRs. Additionally, variations in DNA methylation levels do not entirely explain the expression advantages of homologous genes in F_1_ hybrids. We further generated *Bnadcl3^CR^* and *Bnardr2^CR^* loss-of-function mutants to determine whether RNA-mediated DNA methylation (RdDM) selectively affects the subgenome in allopolyploids. Our findings reveal that genomic introgression influences subgenome asymmetry and gene expression, highlighting the benefits of interspecific characteristics.

## RESULTS

### Unbalance expression of subgenomes in F_1_ hybrids

Significant alterations in gene expression, termed “transcriptome shock”, are caused by subgenomic recombination and are commonly observed following interspecific crosses in plants (Quan et al. 2022; Zhang et al. 2021a). In these cases, homologous genes in the dominant subgenome often show higher expression levels than their homologous counterparts in the recessive subgenome. To investigate subgenomic asymmetric expression resulting from interspecific hybridization between *B. rapa* and *B. napus*, two allotriploid *Brassica* species hybrids were generated by crossing two *B. napus* inbred lines (*s70* and *yu25*, A_n_A_n_C_n_C_n_) with a *B. rapa* line (Hort, A_r_A_r_), resulting in the hybrids Hybrid-sh (*s70* × Hort, A_r_A_n_C_n_) and Hybrid-yh (*yu25* × Hort, A_r_A_n_C_n_), respectively (Fig. S1). The acetocarmine staining assay results showed that the maternal line (*s70* and *yu25*), paternal line (Hort), and F_1_ hybrids contained 38, 20, and 29 chromosomes, respectively (Fig. S1). Although the two *B. napus* inbred lines look similar, the phenotypes of Hybrid-sh were closer to those of the maternal parent, while the phenotypes of Hybrid-yh exhibited more remarkable similarity to the paternal parent, particularly in terms of epidermal color accumulation (Fig. S1). Consistent with phenotypic coloration, the highest accumulation of anthocyanins was observed in the paternal, followed by Hybrid-sh, Hybrid-yh, and the lowest in the maternal (Fig. S2).

The RNA-seq, ATAC-seq, WGBS, and small RNA-seq were then conducted to investigate the regulation of gene expression following allotriploid hybridization (Quan et al. 2024). To accurately compare the transcriptional differences between the parent plants and their F_1_ hybrids across the various subgenomes, the independent *in silico* “hybrids” were created. These hybrids were constructed by combining the sequencing data of parental individuals in a ratio of 1:1 for *B. rapa* and *B. napus*, resulting in the creation of in *silico-sh* (*s70* + Hort, A_r_A_n_C_n_) and *in silico*-yh (*yu25* + Hort, A_r_A_n_C_n_), respectively (see Material and method). Sequencing analysis involved mapping the RNA-seq, ATAC-seq, WGBS, and small RNA-seq reads to a synthetic genome based on the reference genomes of *B. rapa* and *B. napus*, retaining only uniquely mapped reads. Spearman’s rank correlation coefficients confirmed the high quality of these datasets (Tables S1-S4). Consequently, 19,481 homologous triplet genes were identified in the synthetic genomes of *B. rapa* and *B. napus*, with each triplet gene demonstrating a 1:1:1 correspondence across the A_r_, A_n_, and C_n_ subgenomes.

The expression of all subgenome genes, homologous genes, and subgenome-unique genes was compared in F_1_ hybrids and *in silico* ‘hybrids’. The results demonstrated that F_1_ hybrids exhibited the highest expression of A_n_ subgenome genes among all genes and unique genes, followed by C_n_ and A_r_ (Fig. 1a). There was no difference in the expression levels of homologous genes between A_n_ and C_n_ subgenomes; however, the expression of homologous genes in both the A_n_ and C_n_ subgenomes was significantly higher than that in the A_r_ subgenome (Fig. 1a). These observations were also confirmed in other *Brassica* allotriploids (Fig. S3a) (Quan et al. 2022). The distinct expression patterns of genes were then compared between F_1_ hybrids and *in silico* ‘hybrids’, focusing on both unique and homologous genes. In unique genes, no significant differences were found in the expression of the A_r_ and C_n_ subgenomes in *in silico* ‘hybrids’ (Fig. 1b). However, the F_1_ hybrids demonstrated significantly higher expression in the C_n_ subgenome compared to the A_r_ subgenome (Fig. 1a). When examining homologous genes, there were no significant differences in the expression of the A_n_ and C_n_ subgenomes between the F_1_ hybrids (Fig. 1a). In contrast, the *in silico* ‘hybrids’ showed that the expression of the A_n_ subgenome was significantly higher than that of C_n_ (Fig. 1b). These findings indicate that the gene expression patterns following the fusion of distinct subgenomes were markedly different from those predicted by the *in silico* ‘hybrids’. Compared to the simulated values, the expression of the C_n_ subgenome was considerably underestimated after genome recombination.

**Figure 1.**
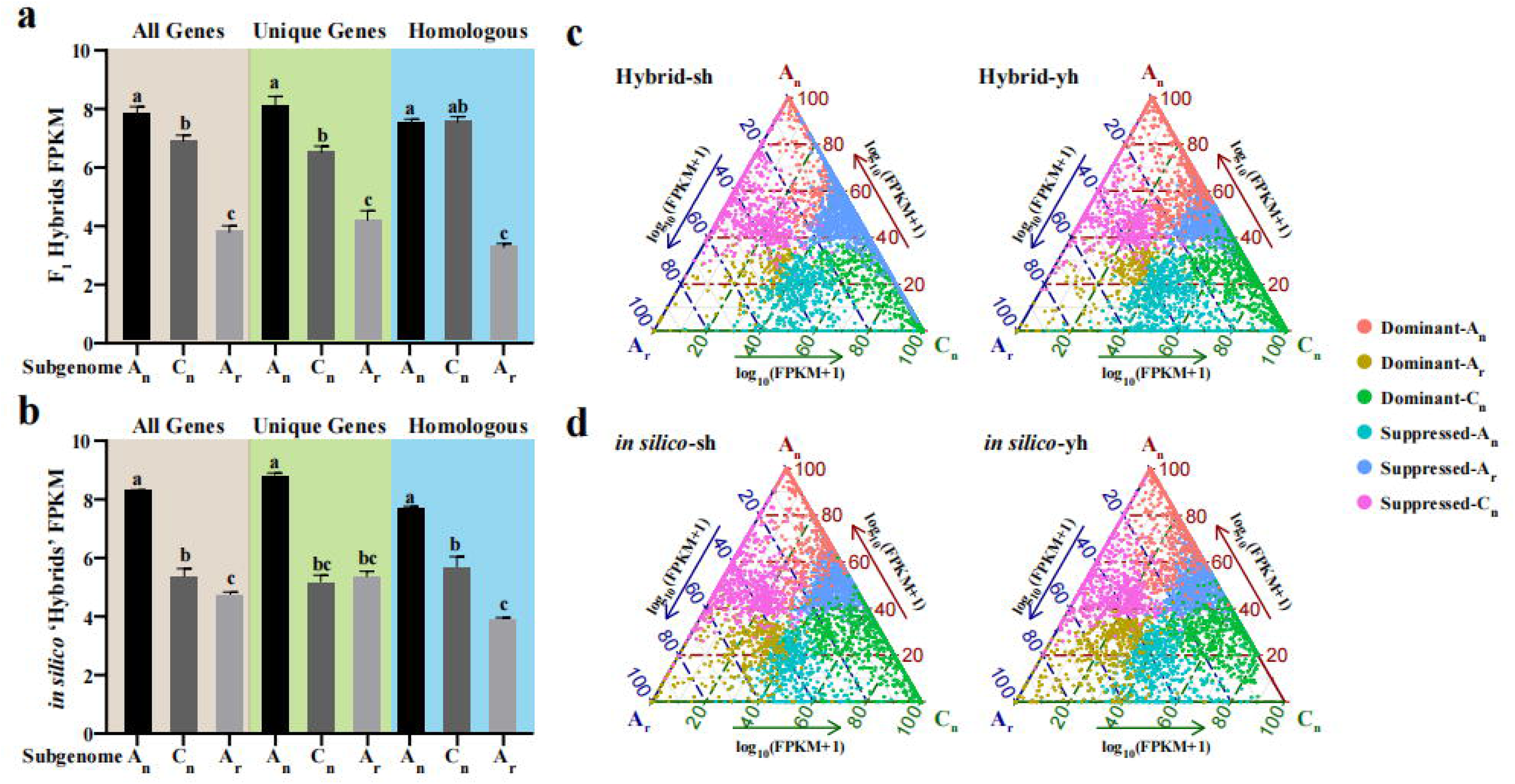
Differentially expressed genes between the three subgenomes in F_1_ hybrids and *in silico* ‘hybrids’. **(a-b)** Histograms showing the expression levels (fragments per kilobase per million, FPKM) of all genes, subgenomic unique genes, and homologous genes in the A_n_, C_n_, and A_r_ subgenomes in the two hybrids **(a)** or two *in silico* ‘hybrids’ **(b)**. **(c-d)** Ternary plots showing expression bias of homologous triplet genes in three subgenomes in F_1_ hybrids **(c)** or *in silico* ‘hybrids’ **(d)**. A_n_-dominant, C_n_-dominant, and A_r_-dominant: higher expression from A_n_, C_n_, or A_r_ subgenomes than the other two homologous genes. A_n_-suppressed, C_n_-suppressed, and A_r_-suppressed: lower expression from A_n_, C_n_, or A_r_ subgenomes than the other two homologous genes.

To investigate how genome fusion affects transcriptome re-editing during interspecific hybridization, a ternary plot was used to visualize the expression bias of homologous triplet genes. We identified the dominant and suppressed homologous triplet genes by comparing their expression levels across the A_n_, A_r_, and C_n_ subgenomes in F_1_ hybrids, applying the criteria of an adjusted *p*-value < 0.05 and a |log_2_ fold change| ≥ 1.5. For instance, if the expression of a homologous triplet gene is higher in the A_n_ subgenome than in both the A_r_ and C_n_ subgenomes (A_n_ > A_r_ and C_n_), we classify it as an A_n_ subgenomic dominant gene. Conversely, if the expression in the A_n_ subgenome is lower than in the A_r_ and C_n_ subgenomes (A_n_ < A_r_ and C_n_), it is categorized as an A_n_ subgenomic suppressed gene. The results from the interspecific hybridization revealed significant changes in the expression of thousands of homologous triplet genes across the subgenomes. Specifically, 3,347 and 3,888 triplet genes showed repression, while 1,927 and 2,350 triplet genes were dominant in Hybrid-sh and Hybrid-yh, respectively (Fig. 1c and Fig. S3b). This indicates that triplet genes in Hybrid-yh are more imbalanced, exhibiting greater suppression than dominance than F_1_ hybrids. Additionally, we identified 712, 1,408, and 2,157 dominant homologous triplet genes in the A_r_, A_n_, and C_n_ subgenomes, respectively. The number of suppressed homologous triplet genes in the A_r_, A_n_, and C_n_ subgenomes was 4,190, 1,132, and 1,193, respectively (Fig. 1c and Fig. S3b). These findings suggest that the C_n_ subgenome exhibits dominance in the expression of homologous triplet genes among the three subgenomes, while the A_r_ subgenome shows suppression. Notably, the F_1_ hybrids displayed more dominant triplet genes and fewer suppressed triplet genes than the *in silico* ‘hybrids’ (Fig. 1d and Fig. S3b). This indicates that many genes were activated rather than repressed during the allotriploid hybridization process.

### The distribution of ACRs in the subgenomes exhibited divergence in F_1_ hybrids

To investigate the dynamics of the accessible chromatin landscape in each subgenome of the allopolyploid, we constructed genome-wide maps of accessible chromatin regions (ACRs) for both F_1_ hybrids and *in silico* hybrids. The Spearman’s rank correlation coefficients generated from the ATAC-seq libraries showed strong reproducibility across biological replicates (Table S3). A principal component analysis (PCA) was conducted to compare the enrichment values at all ACRs across the samples. As previously reported, the biological replicates clustered closely (Quan et al. 2024). Following the criteria established by Tian et al. (2021), overlapping peaks were designated as ACRs if present in at least two of the three biological replicates of each parent and F_1_ hybrid. As a result, we identified 72,947, 61,316, 31,046, 35,639, and 38,553 ACRs in *s70*, *yu25*, Hybrid-sh, Hybrid-yh, and Hort, respectively (Table S5). In both F_1_ hybrids and *in silico* ‘hybrids’, the density of ACRs in the C_n_ subgenome was consistently much higher than in the A_n_ and A_r_ subgenomes (Fig. S4a). These results suggest that the variations in the number of ACRs across different subgenomes in hybrids may indicate subgenome-specific regulatory differences and interspecific hybridization may lead to the silencing or activation of specific genomic regions, resulting in variations in ACRs across the different subgenomes.

The distribution of ACRs across the genome was categorized based on their proximity to genes. They were classified as genic (overlapping with a gene), proximal (within 2 kb of a gene), and distal (more than 2 kb away from any gene). Distinct ACRs in these three categories were observed in F_1_ hybrids between A_n_, A_r_, and C_n_, especially in the proximal (41.6% in A_n_ vs. 47.0% in A_r_ vs. 37.3% in C_n_) and distal (37.2% in A_n_ vs. 29.2% in A_r_ vs. 45.8% in C_n_) regions (Fig. 2a). Compared to the parental lines, the number of distal ACRs in the A_n_ and C_n_ subgenomes increased in the F_1_ hybrids (Fig. 2a). In contrast, the number of proximal ACRs of A_n_ and C_n_ subgenomes was decreased in the two hybrids (Fig. 2a). Notably, the reduction in genic ACRs was matched by an increase in proximal ACRs in the A_r_ subgenome (Fig. 2a). This indicates that ACR adjacent to genes in the A_n_ and C_n_ subgenomes, as well as genic ACRs in the A_r_ subgenome, were suppressed following genome recombination. Conversely, the distal ACRs in the A_n_ and C_n_ subgenomes and proximal ACRs in the A_r_ subgenome were activated.

**Figure 2.**
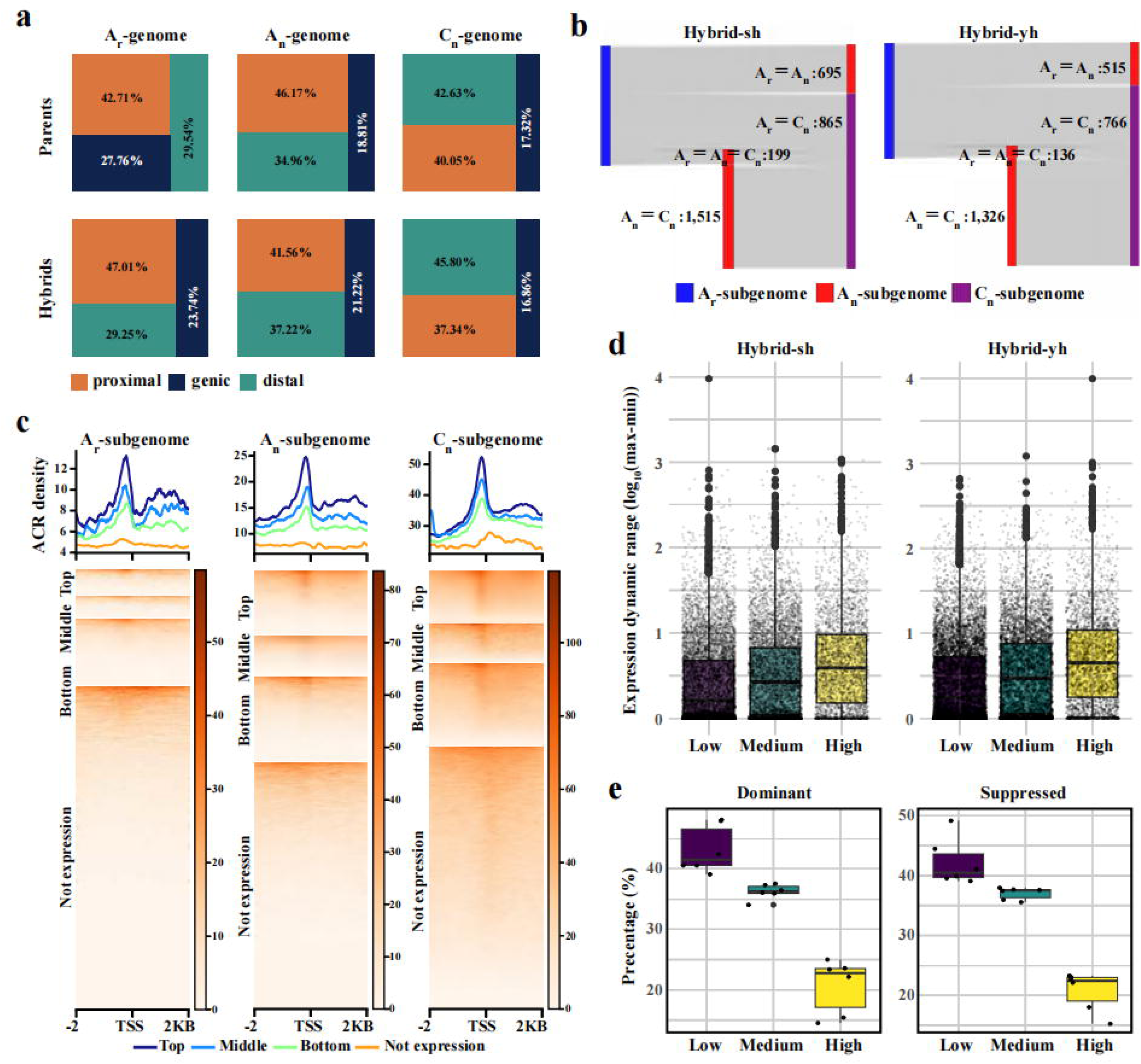
Distribution of subgenomic accessible chromatin regions (ACRs) and correlation with gene expression in F_1_ hybrids. **(a)** Graph showing the proportion of ACRs in genic (overlapping with a gene), proximal (within 2 kb of gene), and distal (more than 2 kb away from any gene) in F_1_ hybrids subgenomes and their parental lines subgenomes. **(b)** Sankey plot showing the comparison of the number of ACRs in homologous triplet genes. **(c)** ACRs around gene transcription start sites (TSS) in the A_n_, A_r,_ and C_n_ subgenomes. Expressed genes were divided into four groups according to FPKM values: Top expressed (FPKM ≥ 10), Middle expressed (5 ≤ FPKM <10), Bottom expressed (1 ≤ FPKM < 5), and not expressed (FPKM ≤ 0.1). Metaplots above each heatmap are derived from genes binned by expression levels: top, middle, bottom, and not expressed. **(d)** Correlation between ACRs regulatory complexity and gene expression variation in F_1_ hybrids. Boxplots showing the distribution of dynamic expression ranges of genes in groups with different complexities. The complexity was divided into three levels according to the number of ACRs: low (0 - 1 ACR), medium (2 - 3 ACRs), and high (≥ 4 ACRs). **(e)** Percentage of low-, medium- and high-complexity genes in dominant and suppressed subgenomes.

To further explore the distribution of ACRs in the A_n_, A_r_, and C_n_ subgenomes after hybridization, we examined the presence of ACRs in homologous triplet genes in two F_1_ hybrids. 14,119 and 14,113 triplet genes contained at least one proximal ACR in Hybrid-sh and Hybrid-yh, respectively. Additionally, 3,163 triplet genes contained no ACR in either Hybrid-sh or Hybrid-yh. We then analyzed whether the A_n_, A_r_, and C_n_ subgenomes have the same number of ACRs in F_1_ hybrids. The results revealed that 51.7-56.1% of homologous triplet genes showed a pattern with more C_n_ ACRs than A_n_ and A_r_ (Fig. S4b). We categorized these homologous triplet genes into four patterns based on their ACR numbers: A_r_ = A_n_ ≠ C_n_, A_r_ = C_n_ ≠ A_n_, A_n_ = C_n_ ≠ A_r_, and A_r_ = A_n_ = C_n_ (Fig. S4c). The pattern A_n_ = C_n_ ≠ A_r_ was the most common, followed by A_r_ = C_n_ ≠ A_n_ and A_r_ = A_n_ ≠ C_n_. In contrast, the pattern A_r_ = A_n_ = C_n_ was the least frequent (Fig. 2b). It appears that A_n_ and C_n_ homologous genes have similar numbers of ACRs, which may explain why there is no significant difference in the expression levels of these homologous genes (Fig. 1a).

### The expression of homologous triple genes in dominant and suppressed subgenomes is associated with chromatin accessibility in F_1_ hybrids

To provide a comprehensive understanding of the intricacies of chromatin accessibility and the regulation of gene expression. We categorized gene expression into four levels: top expressed (FPKM > 10), middle expressed (5 < FPKM < 10), bottom expressed (1 < FPKM < 5), and not expressed (FPKM < 0.1). Our findings indicated that, compared to genes that were not expressed, the top-expressed, middle-expressed, and bottom-expressed genes showed characteristic ACR peaks at the transcription start site (TSS) (Fig. 2c). The density of ACR was highest in top-expressed genes, followed by middle-expressed and then bottom-expressed genes (Fig. 2c). This suggests a strong positive correlation between chromatin accessibility and the expression levels of neighboring genes (R = 0.86). Interestingly, the C_n_ subgenome appears to require greater chromatin accessibility to achieve expression levels comparable to those of the A_n_ and A_r_ subgenomes (Fig. 2c). For example, the ACR densities of middle-expressed genes in the A_r_, A_n_, and C_n_ subgenomes were 10.2, 18.6, and 45.5, respectively (Fig. 2c).

To better understand the complexities involved in homologous triplet gene regulation, we quantified the number of gene-proximal region ACRs and defined regulatory site complexity as the total number of assigned gene ACRs. The genes were then categorized into three groups based on their regulatory complexity: low-complexity genes (no ACR or one ACR), medium-complexity genes (two to three ACRs), and high-complexity genes (four or more ACRs) (Fig. 2d). A significant increase in the dynamic range of gene expression for triplet genes was identified across low, medium, and high complexity levels (Fig. 2d). Additionally, an analysis of the relationship between the dominant and suppressed subgenomes and gene regulatory complexity indicated that the biased expression of triplet genes was more closely associated with low complexity compared to medium or high complexity (Fig. 2e). This suggests that the expression of homologous genes in the dominant and suppressed subgenomes is more closely related to the presence or absence of ACRs, although the link between the number of ACRs and gene expression was not as clear.

### The expression of homologous triple genes in dominant and suppressed subgenomes is not directly associated with DNA methylation in F_1_ hybrids

DNA methylation is a crucial epigenetic modification that plays a significant role in gene regulation and maintaining genome stability (Law et al., 2010). To explore the connection between DNA methylation and the dynamics of subgenome dominance and suppression in allopolyploids resulting from interspecific hybridization, we employed whole-genome bisulfite sequencing (WGBS) to evaluate the levels of DNA methylation (CG, CHG, and CHH) in the promoters and gene bodies of subgenome genes, homologous genes, and subgenome-unique genes in F_1_ hybrids. Across all gene contexts in F_1_ hybrids, the C_n_ subgenome consistently exhibited higher methylation levels compared to the A_n_ and A_r_ subgenomes (Fig. 3a). In homologous gene pairs, both the A_n_ and C_n_ subgenomes showed similar levels of DNA methylation across the gene bodies, which were notably higher than those observed in the A_r_ subgenome in F_1_ hybrids (Fig. 3a). Additionally, the C_n_ subgenome demonstrated greater methylation levels in the proximal region of the gene body compared to homologous genes in the A_n_ and A_r_ subgenomes (Fig. 3a). Interestingly, the DNA methylation level of subgenome-unique genes surpassed that of all genes and homologous genes combined (Fig. 3a). The methylation pattern of these unique genes closely resembled that of all genes, suggesting that subgenome-unique genes primarily drive the DNA methylation levels within subgenomes.

**Figure 3.**
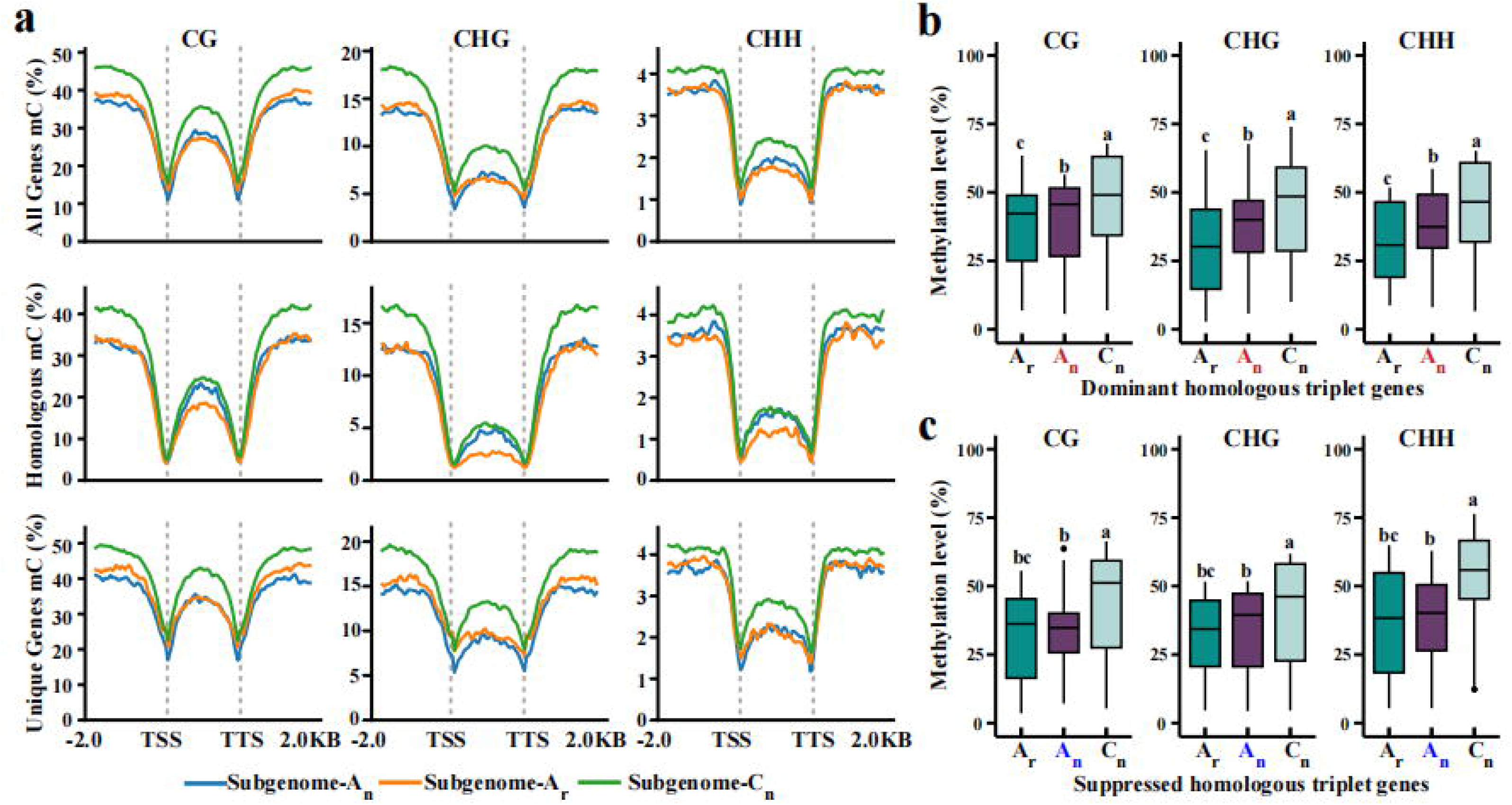
DNA methylation landscape of different subgenomes in F_1_ hybrids. **(a)** DNA methylation levels of all genes, subgenome-unique genes and homologous triplet genes in the A_n_, A_r_, and C_n_ subgenomes. **(b-c)** DNA methylation levels of dominant subgenomic homologous triplet genes **(b)** and suppressed subgenomic homologous triplet genes **(c)** in the A_n_, A_r_, and C_n_ subgenomes. In **(b)** and **(c),** red and blue letters indicate the dominant and suppressive subgenome in the homologous triplet genes, respectively. Average DNA methylation levels were determined from two biological replicates per genotype.

In F_1_ hybrids, the levels of TE (transposable element) bodies methylation in CG and CHG contexts were lower in the A_n_ and A_r_ subgenomes compared to the C_n_ subgenome (Fig. S5a). However, there were no differences in TE body methylation levels between the A_n_ and A_r_ subgenomes (Fig. S5a). In contrast, CHH methylation levels in the A_r_ and C_n_ subgenomes remained unchanged but were higher than in the A_n_ subgenome (Fig. S5a). TEs were categorized into two groups: long TEs (length > 500 bp) and short TEs (length < 500 bp) (Li et al. 2022). For long TE bodies, there were no significant differences in the methylation levels among the A_n_, A_r_, and C_n_ subgenomes in CG and CHG contexts (Fig. S5b). Nevertheless, in the CHH context, the A_n_ subgenome exhibited the lowest methylation level compared to the A_r_ and C_n_ subgenomes (Fig. S5b). In short TE bodies within the CHH context, the A_n_ subgenome showed the highest methylation level, followed by C_n_, while A_r_ had the lowest methylation level (Fig. S5c). These findings indicate that the A_n_ subgenome regulates TE methylation differently from the A_r_ and C_n_ subgenomes, particularly at CHH sites.

We then hypothesize that the expression of dominant and suppressed homologous triplet genes may be directly associated with DNA methylation in the promoter region. The DNA methylation levels of these genes in F_1_ hybrids were quantified. In dominant homologous triplet genes, the methylation levels were highest in the C_n_ subgenome, followed by those in the A_n_ and A_r_ subgenomes (Fig. 3b). The same highest DNA methylation level was also found in C_n_-suppressed homologous triplet genes (Fig. 3c). However, no significant difference in DNA methylation was found between the A_n_, and A_r_-suppressed homologous triplet genes (Fig. 3c). In the gene body region, the A_n_-suppressed homologous triplet genes exhibited higher methylation levels than both the C_n_ and A_r_ subgenomes; however, this trend was not observed in the A_n_ dominant subgenome homologous genes (Fig. S6a and S6b). These findings suggest that the expression of homologous triplet genes, whether dominant or suppressed, is not directly associated with DNA methylation.

### Reduced CHH methylation in F_1_ hybrids was related to DNA methyltransferases and RdDM pathway

Genome-wide DNA methylation is often reprogrammed due to “epigenetic shock” from different parental epigenomes. The average DNA methylation levels of F_1_ hybrids and *in silico* ‘hybrids’ were analyzed. CHH methylation in each subgenome of F_1_ hybrids was reduced compared to *in silico* ‘hybrids’ at the chromosomal level, while the same regions showed unaltered CG and CHG methylation levels (Fig. S7a). The regulation of DNA methylation status relies on three interrelated pathways: DNA methyltransferases, DNA demethylases, and RNA-mediated DNA methylation (RdDM) (Yang et al., 2016). To investigate the causes of reduced CHH methylation in F_1_ hybrids, genes associated with DNA methyltransferases, DNA demethylases, and the RdDM pathway were annotated in the reference genome of *B. rapa-B. napus*. The expression levels of these genes were assessed in both F_1_ hybrids and *in silico* ‘hybrids’. 21 orthologous methyltransferase genes were identified, including *DOMAINS REARRANGED METHYLASE2* (*DMR2*) (6 genes), *CHROMOMETHYLASE 2* (*CMT2*) (3 genes), *CMT3* (3 genes), and *METHYLTRANSFERASE1* (*MET1*) (9 genes). A phylogenetic tree was subsequently constructed (Fig. S7b). Among these 21 genes, *MET1* and *CMT3*, responsible for maintaining CG and CHG methylation, showed minimal or no expression (Fig. S7b). CHH methylation is regulated by two distinct DNA methyltransferases: *CMT2* and *DRM2* (Stroud et al., 2014; Matzke et al., 2014). Compared to the *in silico*-yh, only one of the three orthologous genes for *CMT2* showed decreased expression in the Hybrid-yh samples, while the other two remained unchanged (Fig. S8b). Additionally, the expression levels of all six *DRM2* orthologous genes in Hybrid-yh were lower than those *in silico*-yh (Fig. S7b).

To further investigate whether the RdDM pathway is attenuated following interspecific hybridization, we analyzed the expression levels of several key genes involved in this pathway, including *RNA polymerase IV* (*Pol IV*), *RNA-dependent RNA polymerase 2* (*RDR2*), *Dicer-like 3* (*DCL3*), *RNA polymerase V* (*Pol V*), *Argonaute proteins 4 and 6* (*AGO4/6*), and *DRM2* (Fig. 4a). Compared to *in silico*-yh, A reduction in expression was observed for *BnaA06.DCL3*, *BraA06.DCL3*, *BnaA04.AGO4*, *BraA04.AGO4*, and six *DRM2* orthologous genes (Fig. 4a). To obtain a more direct assessment of RdDM pathway, the abundance of siRNAs and siRNA clusters was quantified and characterized in both F_1_ hybrids and *in silico* ‘hybrids’. The accumulation of siRNAs in the three subgenomes of the F_1_ hybrid showed reduction compared to the *in silico* hybrids, with the A_r_ subgenome exhibiting the most significant decrease, followed by the A_n_ and C_n_ subgenomes (Fig. 4b). A total of 52,651 and 50,082 24-nt siRNA clusters were identified in *in silico*-yh and Hybrid-yh, respectively (Fig. S7c). Compared to *in silico*-yh, we observed a decrease in 24-nt siRNA clusters primarily in the A_r_ subgenome (6565) and the A_n_ subgenome (3391). Conversely, 24-nt siRNA clusters in the C_n_ subgenome increased (7529) (Fig. S7c). This indicates that the reduction of 24-nt siRNA clusters primarily occurs in the A (A_n_ and A_r_) subgenome. In addition, 642 hyper-differential small RNA regions (hyper-DSRs) and 4,455 hypo-differential small RNA regions (hypo-DSRs) were identified in the Hybrid-yh compared to the 24-nt siRNA clusters of *in silico-yh* (Fig. 4c). We also examined the effects of hybridization on DNA methylation levels. Our findings revealed that CG methylation levels remained unchanged (Fig. 4d). In contrast, CHG and CHH methylation levels declined (Fig. 4d). The reduction in RdDM-mediated DNA methylation following hybridization is primarily due to changes in the non-CG methylation context. Furthermore, we found that the long TEs exhibited elevated CG and CHG methylation levels compared to short TEs (Fig. 4e). Conversely, short TEs were identified as a primary contributor to CHH methylation (Fig. 4e).

**Figure 4.**
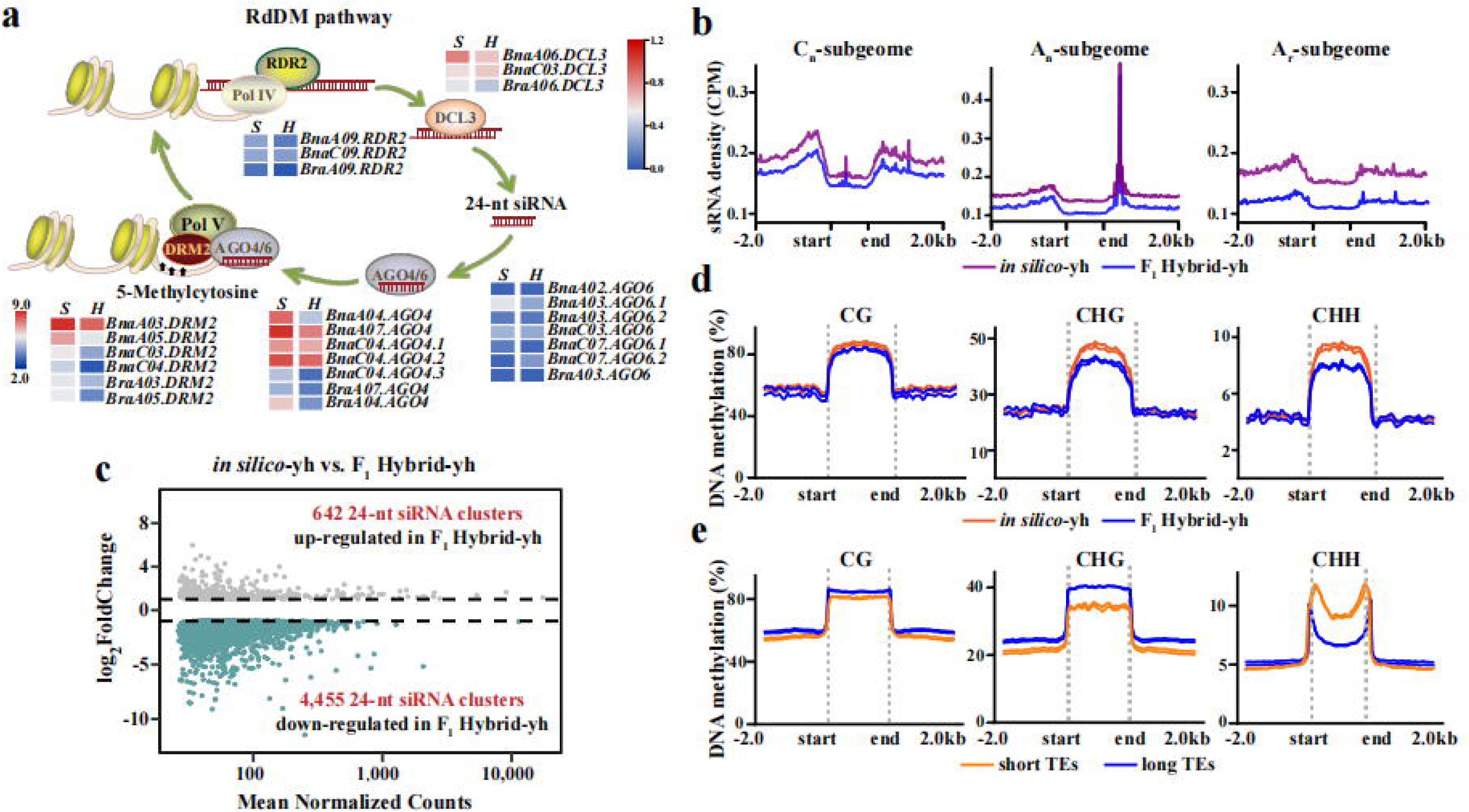
Relationship between the expression of DNA methylation-related genes and CHH hypo-methylation. **(a)** A brief working model of the RdDM pathway showing the transcription levels of genes involved in the RdDM pathway in the Hybrid-yh and *in silico*-yh. *RNA-Dependent RNA Polymerase 2* (*RDR2*) and *Dicer-like proteins* (*DCLs*) are essential for the biogenesis of small interfering RNAs (siRNAs), while *Argonaute proteins 4 and 6* (*AGO4/6*) and *DNA methyltransferase 2* (*DRM2*) participate in siRNA-guided DNA methylation, Heatmap showing transcription levels of these genes involved in RdDM pathways. *H* and *S* represent Hybrid-yh and *in silico*-yh, respectively. **(b)** Distribution of A_n_, A_r_ and C_n_ subgenomic small RNAs in the Hybrid-yh and *in silico*-yh (CPM, counts per million). **(c)** MA-plot showing the number of differentially expressed 24-nt siRNA clusters in the Hybrid-yh compared to *in silico*-yh. The gray and blue dots represent up-regulation and down-regulation, respectively. **(d)** DNA methylation profiles around the Hybrid-yh and *in silico*-yh genes. **(e)** DNA methylation profiles around long TEs and short TEs.

### *BnaDCL3* and *BnaRDR2* maintained non-CG methylation differentially in the A_n_ and C_n_ subgenomes

DNA methylation protects genome stability and ensures the silencing of transposable elements, which helps prevent transposition from destabilizing the host genome (Law et al., 2010; He et al., 2011). To explore whether RNA-mediated DNA methylation occurs preferentially in the non-CG context, we generated *Bnadcl3^CR^* and *Bnardr2^CR^* mutants in *B. napus*. We confirmed deletions or insertions at the sgRNA target sites through sequencing. In the *Bnadcl3^CR^* mutant, the *BnaA06.DCL3* exhibited single base insertions and a 4-bp deletion at two sgRNA target sites, while the *BnaC03.DCL3* showed a single base insertion at the two sgRNA target sites (Table S6). In the *Bnardr2^CR^* mutant, we detected a single base insertion and a 30-bp deletion in *BnaA09.RDR2*, while the *BnaC09.RDR2* presented a 5-bp deletion and a single base insertion (Table S6).

The *Bnadcl3^CR^*, *Bnardr2^CR^*, and wild-type (WT) samples were then subjected to whole-genome bisulfite sequencing (WGBS), achieving approximately 30-fold genome coverage with bisulfite conversion rates exceeding 99% across all samples (Table S7). Spearman’s rank correlation coefficients indicated good reproducibility among biological replicates (Table S7). When comparing the mutants to the wild type (WT), we found that the total number of methylated cytosines was reduced in the CG, CHG, and CHH contexts by 13.7%, 18.5%, and 21.0%, respectively, in *Bnadcl3^CR^*. In *Bnardr2^CR^*, these reductions were 4.0%, 9.2%, and 29.6%, respectively. The entire genome was then segmented into 200 bp windows, which showed similar patterns of methylation levels (Fig. 5a). The most significant reductions in methylation levels for both *Bnadcl3^CR^* and *Bnardr2^CR^* were primarily observed in the CHG and CHH contexts (Fig. 5a). We identified differentially methylated regions (DMRs) between the *Bnadcl3^CR^* and *Bnardr2^CR^* mutants and the WT, discovering 4,929 hypo-DMRs in the CHG context and 11,622 hypo-DMRs in the CHH context for *Bnadcl3^CR^* (Fig. 5b). Notabaly, the *Bnardr2^CR^* mutant displayed a significant increase in these numbers, with 10,950 hypo-DMRs in the CHG context and 46,701 hypo-DMRs in the CHH context (Fig. 5c). Consistent with our observations of differentially methylated regions, we noted that the methylation levels for *Bnadcl3^CR^* and *Bnardr2^CR^* decreased by 40% and 47.1% in the CHH context, respectively (Fig. 5a). Interestingly, in the CHG context, the methylation level for *Bnadcl3^CR^* decreased by 38.2%, which was greater than the 21.5% decrease observed for *Bnardr2^CR^* (Fig. 5a). This indicates that CHH methylation is more dependent on *Bnardr2^CR^* than on *Bnadcl3^CR^*.

**Figure 5.**
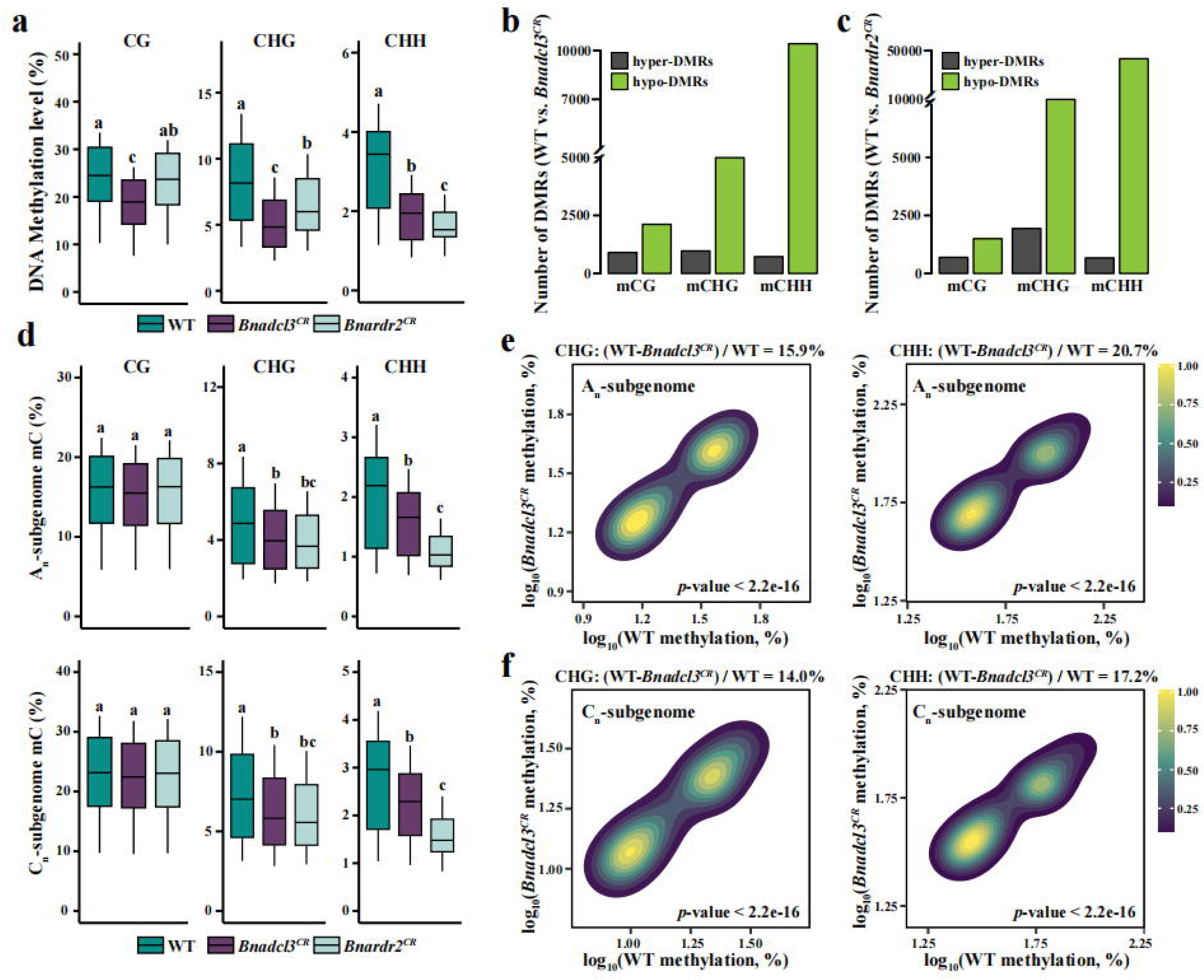
DNA methylation patterns of *Bnadcl3^CR^* and *Bnardr2^CR^* mutants in the A_n_ and C_n_ subgenomes of *B. napus*. **(a)** Boxplots showing the three methylation contents (CG, CHG, and CHH) of *Bnadcl3^CR^* and *Bnardr2^CR^* mutants and the corresponding wild type (WT). Average DNA methylation levels were determined from two biological replicates per genotype. **(b-c)** The number of Hypo- and hyper-DMRs found in the three methylation contexts of *Bnadcl3^CR^* **(b)** and *Bnardr2^CR^* mutants **(c)**. **(d)** Boxplots showing the methylation level of WT, *Bnadcl3^CR^,* and *Bnardr2^CR^* mutants in the A_n_ subgenome (top) and C_n_ subgenome (bottom). **(e-f)** Pairwise comparison of CHG and CHH methylation in WT and *Bnadcl3^CR^*mutants in the A_n_ subgenome **(e)** and C_n_ subgenome **(f)**. The color scale measures the density of points.

We then monitored the methylation levels of genes from both the A_n_ and C_n_ subgenomes. The CG methylation levels of *Bnadcl3^CR^* and *Bnardr2^CR^* were not found to be diminished within the gene body (Fig. 5d). However, significant reductions in CHG methylation levels were observed in both *Bnadcl3^CR^* and *Bnardr2^CR^* compared to the wild-type (Fig. 5d). There was no significant difference in the extent of the reduction between *Bnadcl3^CR^* and *Bnardr2^CR^* (Fig. 5d). The greatest decrease in methylation levels was found in *Bnardr2^CR^*, particularly regarding the CHH sequence in both the An and Cn subgenomes (Fig. 5d). In the context of mutant *Bnadcl3^CR^*, the methylation levels decreased by 15.9% in the A_n_ subgenome and by 14.0% in the C_n_ subgenome for CHG, while decreases of 19.2% and 16.7% were observed in the A_n_ and C_n_ subgenomes for CHH, respectively (Fig. 5e and 5f). Similar trends were noted for *Bnardr2^CR^*, with decreases of 19.2% (CHG) and 45% (CHH) in the A_n_ subgenome, and 16.8% (CHG) and 41.4% (CHH) in the C_n_ subgenome (Fig. S8a). These findings suggest that *Bnadcl3^CR^* and *Bnardr2^CR^* have a more significant impact on the DNA methylation levels of the A_n_ subgenome.

The CHH methylation levels of TEs were found to be reduced across the genome in both the *Bnadcl3^CR^* and *Bnardr2^CR^* mutants, with reductions of 13.9% and 21.6%, respectively (Fig. S8b). However, these reductions were less significant than those observed in the gene body and its nearby regions, where reductions were 40.0% for *Bnadcl3^CR^* and 47.1% for *Bnardr2^CR^*. Additionally, the decrease in methylation levels was more significant in short TEs (28.8% for *Bnadcl3^CR^* and 49% for *Bnardr2^CR^*) than in long TEs (13.7% for *Bnadcl3^CR^* and 24.7% for *Bnardr2^CR^*) (Fig. S8c). Consistent with the trends in the gene body, the reduction of CHH methylation in TEs appears to be more closely related to the *Bnardr2^CR^* mutant. These findings suggest that CHH methylation primarily occurs in short TEs.

We then conducted small RNA sequencing (sRNA-seq) on *Bnardr2^CR^*, *Bnadcl3^CR^*, and WT. After removing low-quality reads and structural RNAs, the remaining clean small RNA (sRNA) reads were mapped to the *B. napus* genome (Zhongshuang 11, ZS11) (Song et al. 2020). The abundance of small interfering RNAs (siRNAs) was quantified using reads per million mapped reads (RPM). The sRNA-seq data generated an average of 12 million clean reads per replicate, with 95.8% of these mapped successfully to the reference genome (Table S8). The reproducibility between biological replicates was confirmed using Spearman’s rank correlation coefficients (Table S8). Next, we compared the abundance and genome-wide distribution of siRNAs in *Bnadcl3^CR^*, *Bnardr2^CR^*, and WT. siRNAs were uniformly distributed across the chromosome arms, while there was a noticeable depletion in the pericentromeric region (Fig. 6a). Compared to WT, the accumulation of siRNAs was down-regulated across the entire genome in *Bnadcl3^CR^*, but this decline was less pronounced than in *Bnardr2^CR^* (Fig. 6a). Specifically, in *Bnardr2^CR^*, the accumulation of 22-26 nt siRNAs was significantly reduced, whereas only 24-nt siRNAs showed a significant reduction in *Bnadcl3^CR^* (Fig. 6b). This difference may stem from BnaDCL3’s specific targeting of 24-nt siRNAs, where the quantity of 24-nt siRNAs decreased significantly, while the levels of other types of siRNAs increased (Fig. 6b).

**Figure 6.**
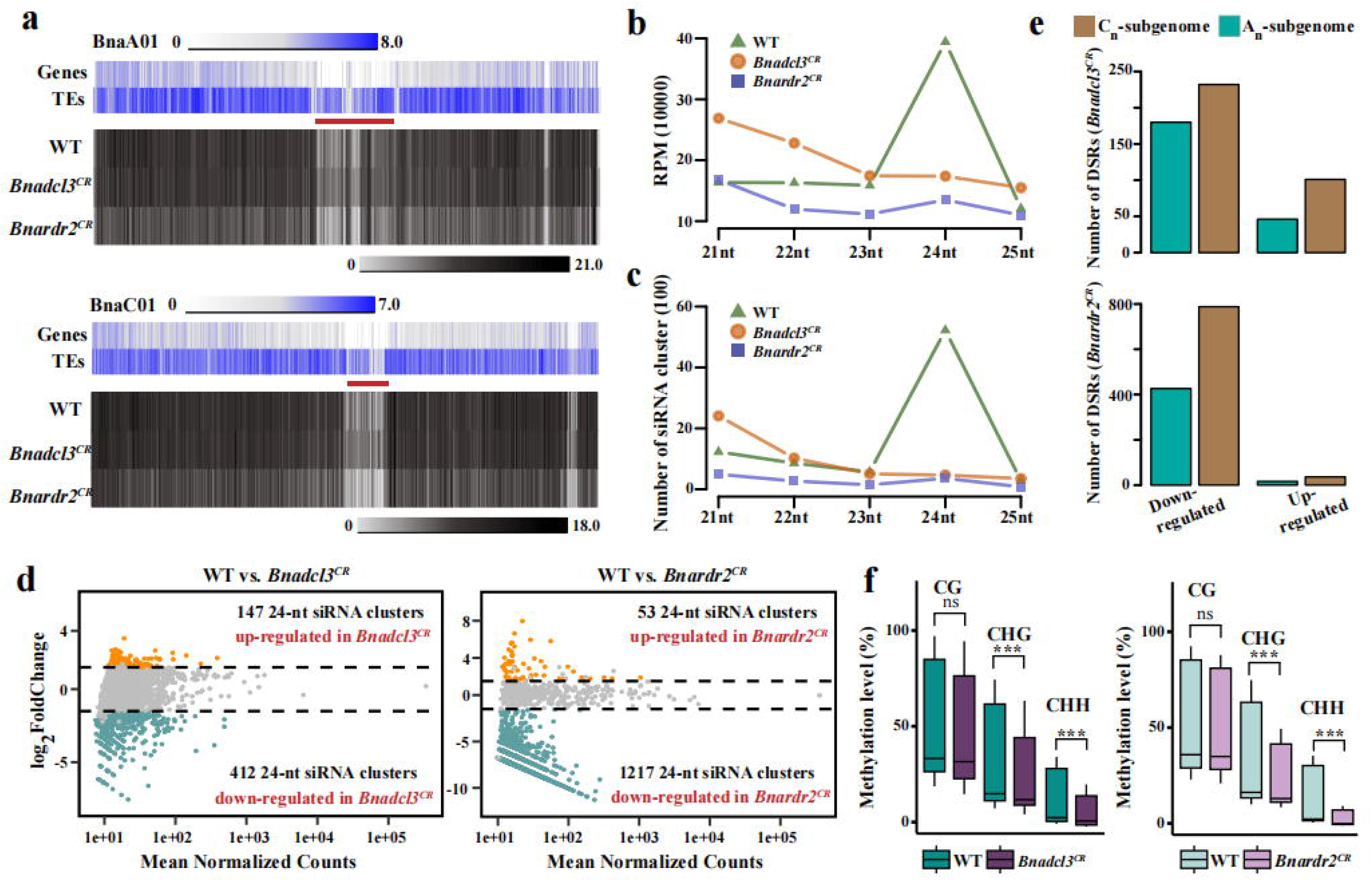
Distribution and differential expression patterns of small RNAs in WT, *Bnadcl3^CR^,* and *Bnardr2^CR^* mutants. **(a)** Whole-genome sRNA heatmap across the genome at 100 kb resolution, top track: gene and TE density. sRNA from WT, *Bnadcl3^CR^* and *BnaRDR2* mutants. The centromere was represented by thick red line. **(b-c)** Graph showing the distribution of siRNA expression levels **(b)** and siRNA clusters **(c)** of WT, *Bnadcl3^CR^*, and *Bnardr2^CR^* mutants. **(d)** The MA-plot showing the number of differentially expressed 24-nt siRNA clusters in *Bnadcl3^CR^* and *Bnardr2^CR^* mutants. The blue and orange dots represent up-regulated and down-regulated, respectively. **(e)** Number of differential small RNA regions (DSRs) in *Bnadcl3^CR^*(top) and *Bnardr2^CR^* (bottom) mutants in the A_n_ subgenome and C_n_ subgenome. **(f)** DNA methylation levels of differentially down-regulated 24-nt siRNA clusters in WT, *Bnadcl3^CR^* (right), and *Bnardr2^CR^* (left) mutants. Error bars indicate mean ± SD from three biological replicates. Student’s *t*-test; \*\*\**p* < 0.001.

Next, we counted the number of 24-nt siRNA clusters and analyzed the differential small RNA regions (DSRs). A total of 5224, 466, and 361 24-nt siRNA clusters were identified in WT, *Bnadcl3^CR^*, and *Bnardr2^CR^* mutants, respectively (see Material and Methods) (Fig. 6c). These results indicate that the synthesis of 24-nt siRNAs in *B. napus* is dependent on *BnaDCL3* and *BnaRDR2*. Notably, the *Bnadcl3^CR^* and *Bnardr2^CR^* mutants retained 4.1% and 2.9% of the 24-nt siRNA clusters from WT plants, respectively (Fig. S9a). When comparing the 24-nt siRNA clusters in WT plants, we identified 147 and 53 hyper-DSRs, as well as 412 and 1,217 hypo-DSRs in the *Bnadcl3^CR^* and *Bnardr2^CR^* mutants, respectively (Fig. 6d). Additionally, most hypo-DSRs were found within short TEs (58.5% in *Bnadcl3^CR^* and 61% in *Bnardr2^CR^*) (Fig. S9b). In the C_n_ subgenome of *Bnardr2^CR^*, the number of hypo-DSRs was 1.9 times higher than that in the A_n_ subgenome, whereas in *Bnadcl3^CR^*, it was only 1.3 times higher (Fig. 6e). These findings suggest that BnaDCL3 and BnaRDR2 exhibit selectivity for the A_n_ and C_n_ subgenomes regarding differential small RNA regions. Hypo-DSRs in the C_n_ subgenome were dependent on BnaRDR2 and were partially influenced by BnaDCL3 (Fig. 6e). Furthermore, hypo-DSRs showed a reduction in CHG and CHH methylation levels in both *Bnadcl3^CR^* and *Bnardr2^CR^* mutants (Fig. 6f). Specifically, in the *Bnardr2^CR^* mutant, CG, CHG, and CHH methylation levels decreased by 3.9%, 29.1%, and 63.1%, respectively (Fig. 6f). In the *Bnadcl3^CR^* mutant, the reductions in methylation were 7.6% for CG, 21.1% for CHG, and 36.1% for CHH (Fig. 6f). These data support the critical roles of *BnaDCL3* and *BnaRDR2* in DNA methylation processes, particularly in maintaining methylation in non-CG contexts.

## DISCUSSION

Interspecific hybridization is a crucial mechanism for the generation of new species and the introduction of phenotypic diversity. However, the asymmetric expression of subgenomes that occurs when multiple genomes merge during interspecific hybridization is still not completely understood. This study aims to clarify the mechanisms underlying the bias in homologous expression and the observed reduction in genome-wide DNA methylation in interspecific hybrids. We employ chromatin accessibility analysis, DNA methylation analysis, and mutants of RdDM pathway genes to investigate the potential factors contributing to the subgenomic advantages.

### Interspecific hybridization affects the distribution of accessible chromatin regions in F_1_ hybrids

Understanding the characteristics of open chromatin, including its distribution throughout the subgenome and its correlation with gene expression, is essential for deciphering the biased expression and dominance of homologous genes derived from different subgenomes. High-resolution maps of open chromatin regions have been generated for species such as *Arabidopsis*, hexaploid wheat, and maize (Tian et al., 2021; Zhao et al., 2023; Yin et al., 2022). We employed ATAC-seq to analyze the distribution of accessible chromatin regions (ACRs) across the genomes of F_1_ hybrids and their parental lines. Our analysis revealed that the number of ACRs in the F_1_ hybrids was significantly lower than in the parental lines. This reduction may result from the recombination and rearrangement of genomes from different species during the formation of allopolyploids (Han et al., 2022; Zhu et al., 2017). Such processes can alter chromatin structure, potentially leading to the loss or closure of certain ACRs following hybridization. Additionally, epigenetic modifications, including DNA methylation and histone modifications, can affect chromatin accessibility (Zhong et al., 2021; Li et al., 2023a). Genome-wide chromatin accessibility maps for 18 *Arabidopsis* mutants that lack CG, CHG, or CHH methylation demonstrated that DNA methylation in all three sequence contexts influences chromatin accessibility (Zhong et al., 2021). When compared to the allotetraploid parental lines, the ACRs in Hybrid-sh and Hybrid-yh decreased by 57.4% and 41.8%, respectively (Table S6). In Hybrid-sh, we observed a significant increase in DNA methylation levels relative to the parental line; however, Hybrid-yh exhibited minimal changes (Quan et al., 2024). This suggests that increased DNA methylation due to hybridization may lead to the silencing of certain ACRs. Consistent with previous findings (Li et al., 2022), the proportions of ACRs in the C_n_ and A_n_ subgenomes of the maternal line were 63.5% and 36.5%, respectively, indicating that the C_n_ subgenome has more accessible chromatin regions. Due to the halving of the copy number of the C_n_ genome in F_1_ hybrids (A_r_A_n_C_n_) compared to the parent (A_n_A_n_C_n_C_n_), some C_n_ subgenome-specific ACRs may be silenced due to gene dosage effects.

### Interspecific hybridization affects the subgenomic bias in allopolyploid F_1_ hybrids

Asymmetric subgenomic gene expression has been revealed in the successful establishment of allopolyploids (e.g., *Brassica*, cotton, tobacco, wheat, maize, sorghum, Medicago, and *Arabidopsis*) (Cheng et al. 2018; Bird et al. 2018). In cotton, 69% of homologous genes exhibited a dynamic bias in expression toward the A_t_ or D_t_ subgenomes across different tissues (Huang et al. 2024). Significant differences in homologous gene expression biases were also identified between the subgenomes in natural *Mimulus peregrinus*, resynthesized *M. peregrinus*, and the interspecific triploid hybrid *M. robertsii* (Edger et al. 2017). The allopolyploid species *B. napus* contains A_n_ and C_n_ subgenomes (Chalhoub et al. 2014). Numerous studies have investigated asymmetric subgenomic dominance in both natural and resynthesized *B. napus*. In cultivars 2063A and B409, and their F_1_ hybrid, the expression levels of genes in the A_n_ subgenome were found to be higher than those in the C_n_ subgenome for both cultivars and hybrids (Zhang et al. 2021b). In contrast, the expression levels of homologous gene pairs did not differ significantly between the A_n_ and C_n_ subgenomes (Zhang et al. 2021b). These findings align with our observations that there were no significant differences in the expression levels of homologous genes between the A_n_ and C_n_ subgenomes in allotriploid F_1_ hybrids. However, other studies of natural *B. napus* (*Darmor*) and resynthesized *B. napus* L. have reported that homologous gene pairs in the C_n_ subgenome tend to dominate over those in the A_n_ subgenome (Bird et al. 2021; Li et al. 2021). These discrepancies suggest that the significant differences in homologous gene expression levels between the A_n_ and C_n_ subgenomes in resynthesized *B. napus* may arise from variations in their parental lines.

In allopolyploids, variations in chromatin accessibility across different subgenomes may significantly influence biased gene expression (Jordan et al. 2020; Li et al. 2022; Yin et al. 2022). By integrating ACRs with transcriptome data analysis, we identified a significant positive correlation between ACRs and gene expression (Fig. 2c), which is consistent with previous studies conducted on hexaploid wheat, *Arabidopsis*, and *B. napus* (Li et al. 2022; Tian et al. 2021; Zhao et al. 2023). When comparing the A_n_ and A_r_ subgenomes, we observed a distinct peak at the transcription start site of the C_n_ subgenome (Fig. S4a). The differences in gene expression among homologous triplet genes are directly related to the presence of ACRs in the regions adjacent to these genes (Fig. 2e). This pattern may help clarify the uneven expression of homologous gene pairs across the subgenomes.

Recent evidence indicates that differences in DNA methylation and histone modifications between subgenomes may contribute to imbalanced subgenome expression (Li et al. 2022; Bird et al. 2021; Li et al. 2021; Zhang et al. 2021b). In resynthesized *B. napus*, the average methylation level of the A_n_ subgenome was found to be lower than that of the C_n_ subgenome (Bird et al. 2021). Furthermore, homologous gene pairs with biased expression in the C_n_ subgenome exhibited higher methylation levels in their flanking regions compared to those in the A_n_ subgenome, while the C_n_ homologous genes demonstrated higher expression than their homologs in the A_n_ subgenome (Zhang et al. 2023). These findings suggest that variations in DNA methylation alone do not fully account for the subgenomic dominance observed in resynthesized *B. napus*. Active histone modification markers, such as H3K4me3 and H3K27ac, were frequently identified in biased-expression homologous genes between the A_n_ and C_n_ subgenomes in *B. napus* (*Darmor*) (Li et al. 2021). Additionally, H3K27me3, a transcriptionally repressive epigenetic mark, was not correlated with biased-expression gene pairs in either resynthesized or natural *B. napus*, consistent with previous findings in rice (Lv et al. 2019). This indicates that active histone modifications are significantly and positively associated with biased gene expression. Genome mismatch and recombination during allopolyploid formation can lead to genome instability (Cao et al. 2023a). To maintain genome stability, plants may inhibit the activity of potentially harmful transposons and repetitive sequences by increasing DNA methylation, particularly in newly integrated or heterologous gene regions (Edger et al. 2017; Zhang et al. 2022). Interestingly, DNA methylation levels in subgenome-specific genes are generally higher than in homologous genes (Fig. 3a). Unique genes may be more susceptible to such regulatory mechanisms because they might have undergone more significant genome recombination and selection pressure throughout evolution. These results suggest DNA methylation surrounding unique genes may be linked to genome stability.

### Reduced CHH methylation is associated with the RDdM pathway

DNA methylation modifications play a significant role in regulating gene expression, maintaining genome stability, and facilitating development (Law et al. 2010; He et al. 2011). Various methylation patterns are managed by different enzymes and pathways. The maintenance of CG and CHG methylation primarily depends on the methyltransferases *MET1* and *CMT3* (Kankel et al. 2003; Lindroth et al. 2001). Since the expression levels of *MET1* and *CMT3* did not change in the F_1_ hybrids (Fig.4a), the methylation levels at CG and CHG sites remained stable in F_1_ hybrids (Fig. S8b). Moreover, DNA demethylases, such as ROS1, DME, and DML2/3, are crucial in regulating the methylation status of the genome (Qian et al. 2012; Lister et al. 2008). These demethylases actively remove methyl groups from DNA, influencing gene expression levels. However, our study did not reveal any significant changes in the activity of these demethylases (Fig. S9c), suggesting their role may not be a major factor in the observed decrease in CHH methylation levels.

We further generated mutants of *RDR2* and *DCL3* in *B. napus*. Our findings revealed that methylation levels for CHG and CHH were reduced, while CG methylation remained unchanged (Fig. 5a). This observation is consistent with previous research conducted on other crops (Wang et al. 2022). In the *Bnadcl3^CR^* and *Bnardr2^CR^* mutants, we observed a more significant reduction in DNA methylation in the A_n_ subgenome compared to the C_n_ subgenome (Fig. 5b). These findings may be related to the composition and relative abundance of the genomes. The density of transposable elements (TEs) in the A_n_ subgenome is higher than in the C_n_ subgenome, and gene expression levels in the A_n_ subgenome also surpass those in the C_n_ subgenome (Zhang et al. 2023; Zhang et al. 2021b). Another contributing factor could be that the C_n_ subgenome exhibits higher levels of ACR and H3K27me3 (Li et al. 2021; Zhang et al. 2021b). Structural and functional differences between the A_n_ and C_n_ subgenomes, such as variations in repeat sequences or the distribution of specific functional regions, may further affect their sensitivity to changes in DNA methylation (Zhang et al. 2023). These results underscore the crucial roles of RDR2 and DCL3 in the plant RNA-directed DNA methylation (RdDM) pathway and highlight their significance in the de novo methylation of CHG and CHH.

## MATERIAL AND METHODS

### Plant material

*Brassica napus* ‘s70’ and ‘yu25’ (*B. napus* L.; AACC, allotetraploid) were chosen as the maternal parents, while *Brassica rapa* species (*B. campestris L. ssp. chinensis var. purpuria* Hort.; AA, diploid) were selected as the paternal parent. The F_1_ allotriploid hybrids were created through a cross between two different *B. napus* and the *B. rapa* species. All plant materials were cultivated under the same field conditions at Huazhong Agricultural University (30°28ʹN, 114°21ʹW). Sampling was conducted between 10.00–11.00 h in December (with an average temperature of 8°C). Stem epidermal tissue from under the fifth true leaf at the top was collected and promptly frozen in liquid nitrogen. These samples were then utilized for subsequent RNA sequencing (RNA-seq), small RNA sequencing (sRNA-seq), assay of transposase accessible chromatin sequencing (ATAC-seq), and whole-genome bisulfite sequencing (WBGS) library construction.

### Vector construction and plant transformation

The *pRGE-BnaDCL3* and *pRGE-BnaRDR2* plant expression vectors were constructed as previously described (Xie et al. 2015). The sgRNAs were selected to target *BnaDCL3* and *BnaRDR2* through CRISPR-P (http://cbi.hzau.edu.cn/cgi-bin/CRISPR). All primers used for vector construction are listed in Supplementary Table 8. *pRGE-BnaDCL3* and *pRGE-BnaRDR2* vectors were introduced into *Agrobacterium rhizogenes* strain *GV3101* and used to genetically transform the Westar background (used as wild type, WT) via the hypocotyl dark-light culture transformation method, as described by Dai et al. (2020).

To analyze the mutations caused by CRISPR/Cas9, genomic DNA was extracted from each transgenic plant using the CTAB method (Molecular Cloning, 3^rd^ edition) and further determined by using the Hi-TOM platform (Liu et al. 2019). Target-specific and barcoding PCR was performed to amplify the genomic region encompassing the specific targets of independent samples, and the resulting PCR products were mixed in equal amounts and purified for next-generation sequencing (Novogene Bioinformatics Institute, China). The resulting sequencing data were then decoded by a corresponding online tool to track the mutations of the target sites (http://www.hi-tom.net/hi-tom/). The target-specific primers were listed in Supplemental Table S9.

*Bnardr2^CR^*, *Bnadcl3^CR^*, and WT (Westar) were cultivated under the same field conditions at Huazhong Agricultural University (30°28ʹN, 114°21ʹW). Sampling was conducted between 10.00-11.00 h in December (with an average temperature of 8°C). Young leaves were collected and quickly frozen in liquid nitrogen. These samples were then utilized for sRNA-seq and WBGS library construction.

### *In silico* hybrids and reference genome construction

We constructed an integrated reference genome based on the alignment result of *B. napus* (*s70* and *yu25*) and *B. rapa* (Hort) re-sequencing and RNA-seq data. First, single nucleotide polymorphisms (SNPs) were called between the *B. napus* (ZS11 v.0)-*B. rapa* (Chiifu v.3.0) reference genome and *B. napus* (*s70* and *yu25*)-*B. rapa* (Hort) re-sequencing alignment using GATK (v.4.0) Unified Genotyper, filtered only to include homozygous SNPs, and the new reference genome was made using GATK_v4.0 FastaAlternativeReferenceMaker (Song et al. 2020; Zhang et al. 2018; McKenna et al. 2010). The *B. napus* (*s70* and *yu25*) reference genome was concatenated to the *B. rapa* (Hort) reference genome to create a reference genome for matching the in silico and F_1_ hybrids in this study.

To ensure accurate comparisons of expression between F_1_ hybrids and their parental lines, we constructed independently *in silico* ‘hybrids’ by combining the RNA-seq, small RNA-seq, ATAC-seq, and WGBS data from sequenced parental individuals in a 1:2 ratio for the *B. napus* (*s70* and *yu25*) and *B. rapa* (Hort) datasets. This ratio reflects the respective 1:2 genomic contribution of the parents to the F_1_ hybrids.

### Identification of dominant or suppressed homologous triplet genes

The OrthoFinder_v2.5.4 was used to identify homologous triplet genes by retrieving the protein sequences of the A_r_, A_n_, and C_n_ subgenomes with default settings (Emms and Kelly, 2015). The dominant or suppressed homologous triplet genes were identified by comparing the expression levels of these genes between the A_n_, A_r_, and C_n_ subgenomes, using the criteria of an adjusted p-value < 0.05 and a |log_2_ fold change| ≥ 1.5. For example, if the expression of a homologous triplet gene is higher in the A_n_ subgenome than in the A_r_ and C_n_ subgenomes (A_n_ > A_r_ and C_n_), it is defined as an A_n_ subgenomic dominant gene. Conversely, if the expression in the A_n_ subgenome is lower than in the A_r_ and C_n_ subgenomes (A_n_ < A_r_ and C_n_), it is classified as an A_n_ subgenomic suppressed gene.

### RNA-seq and small RNA-seq (sRNA-seq) assay

According to the manufacturer’s instructions, total RNA was extracted with the RNeasy Plant Mini Kit (Qiagen, Cat No./ID: 74904). This preparation was split and used for both RNA-seq and sRNA-seq. According to the manufacturer’s protocols, library construction and deep sequencing were performed using the Illumina HiSeq 4000 Platform (Novogene, Beijing, China). Three biological replicates were performed.

For RNA-seq analysis, Trimmomatic_v0.38 (java-jar trimmomatic-0.38.jar PE-threads 4-phred33 TruSeq3-PE.fa:2:30:10 SLIDINGWINDOW:4:15 MINLEN:36 LEADING:3 TRAILING:3) was used to remove barcode adaptors and low-quality reads (Bolger et al. 2014). The filtered reads were then aligned to a synthetic genome of the *B. rapa* and *B. napus* reference genomes using HISAT2_v2.2.0 (hisat2 -p 8 -x) with default parameters (Zhang et al. 2018; Song et al. 2020; Kim et al. 2015). The uniquely mapped reads were filtered using SAMtools_v1.9 (Li et al. 2009). Read counting and normalization of transcripts per million mapped reads (FPKM) were performed on BAM files using StringTie_v2.1.4 (stringtie -e -G -o) (Pertea et al. 2015). Genes with FPKM > 1 were defined as expressed genes. Genes with an adjusted *p*-value < 0.05 found by DEseq2_1.36.0 and a |log_2_fold change| ≥ 1.5 were assigned as differentially expressed (Love et al. 2014). The raw sequencing reads were trimmed using cutadapt to remove adapters (https://github.com/ marcelm/cutadapt/tree/v3.1). Subsequently, sRNAs between 18 and 30 nt in length were selected and mapped to the synthetic genome and defined into sRNA clusters using Shortstack_v3.8.4 (ShortStack--readfile--outdir--genomefile--bowtie_cores 5--sort_mem 10G--mismatches 0--ranmax 5--mincov 1rpm--dicermin 20--dicermax 26) (Johnson et al. 2016). sRNA-mapped reads were normalized to the total cleaned reads for further analysis.

### ATAC-seq analysis

For each biological replicate, the collected plant tissue was cut into small pieces with the blade in 500 mL lysis buffer (15 mM Tris-HCl pH7.5, 20 mM NaCl, 80 mM KCl, 0.5 mM spermidine, 5 mM 2-mercaptoethanol and 0.2% Triton X-100). After confirming nuclear integrity, purified nuclei were resuspended in a 50 μL Tn5 transposase integration reaction and incubated at 37°C for 30 min. The Tn5 transposase-digested DNA fragments were then recovered using a MinElute PCR Purification Kit (Qiagen, Cat No./ID: 28004), followed by purification and amplification. The purified library was then sequenced on an Illumina Novaseq platform by Novogene Gene Technology (Novogene, Beijing, China). All ATAC-seq profiles were generated from at least three independent biological replicates.

The low-quality reads and adapters from the raw ATAC-seq data were filtered and removed using Trimmomatic_v0.38 (java-jar trimmomatic-0.38.jar PE-threads 4-phred33) (Bolger et al. 2014). The clean data were then aligned to the synthetic genome by Bowtie2_v2.5.2 (bowtie2 -p 8 -q -I 10 -X 1000 --dovetail--no-unal--very-sensitive-local--no-mixed--no-discordant -x) (Langmead & Salzberg 2012). The mapped reads in sam format were converted to bam format using SAMtools_v1.9 (samtools view -b -f 2 -q 30 -o) (Li et al. 2009). Subsequently, MACS2_v2.2 (macs2 callpeak -t -n -g --nomodel --shift −100 --extsize 200) peak calling software was used to identify ATAC-seq peaks (Zhang et al. 2008). The overlapping peaks over 50 bp in the biological replicates were considered as ACRs. The genomic distribution of ACRs and associated genes was confirmed using the ChIPseeker_v1.32.1 (Yu et al. 2015). Differential binding events were identified using the DiffBind_v3.6.5 package (Ross-Innes et al. 2012). ACR motifs were identified by using findMotifsGenome.pl in HOMER_v5.1 (Heinz et al. 2010).

### Whole genome bisulfite sequencing assay

Genomic DNA was extracted from 10 samples using the cetyl trimethylammonium bromide (CTAB) method. A lambda DNA spike-in was utilized to correct for non-conversion rates of uracil, with 1 ng of methyl-free lambda DNA added to 1 μg genomic DNA as an internal reference for the conversion test. Bisulfite conversion of DNA was carried out using the EZ DNA Methylation Gold Kit (Zymo Research in Irvine, California, USA). The Bisulfite-Seq Library Prep Kit for Illumina (Novogene in Beijing, China) was used to construct whole genome bisulfite sequencing (WGBS) libraries, which were then sequenced on an Illumina HiSeq X10 platform at a depth of 30-fold. Two biological replicates were performed.

We then used BatMeth2_v.2.01 (BatMeth2 pipel --region 200bp --binCover 4 --Qual 5 −1 −2 -g -p 20 -o --gff) align with default parameters to map the filtered WGBS reads to the synthetic genome (Zhou et al. 2019). The sequences covering five or more cytosine sites were set as valid methylation sites. Finally, BatMeth2-Meth2BigWig was used to generate BigWig files to identify and visualize differentially methylated regions (DMRs) in IGV (BatMeth2 batDMR -g -o_dm -o_dmr −1 −2) (Zhou et al. 2019). Only cytosine regions with adjusted *p*-values < 0.05 and DNA methylation differences greater than 0.3, 0.2, and 0.1 (for CG, CHG, and CHH, respectively) were considered DMRs.

### Quantification of anthocyanins

Total anthocyanins were quantified in stem epidermal tissue as described in previous studies (Gao et al. 2020). Approximately 0.5 g of fresh stem epidermis is ground to a powder using liquid nitrogen. Add 5 mL of extraction solution (methanol: water: formic acid: trifluoroacetic acid in a ratio of 70:27:2:1) and incubate in the dark at 4°C in a refrigerator for 12 hours. Transfer the supernatant by filtration to a new test tube. The absorbance was measured at 530 nm using a UV spectrophotometer (Hoefer Vision, SP-2001). All samples were quantified in triplicate in three independent biological replicates.

### Acetocarmine staining assay

Root tips of F_1_ hybrids and their parental lines were excised and fixed in Carnoy’s fixative buffer (ethanol: acetic acid = 3:1). The fixed root tips were then immersed in a 1:1 mixture of hydrochloric acid and alcohol and dissociated for 30 min at room temperature. After rinsing with distilled water to remove the acid-alcohol mixture, the root tips were placed in an acetocarmine staining solution (LMAI Bio, LM0045, China) for 5 to 10 minutes. Finally, the well-distributed chromosomes were observed under a microscope (Nikon, SMZ25, Japan).

### Phylogenetic tree analysis

The protein sequences of *Brassica* species *DRM2*, *CMT3*, *CMT2*, and *MET1* genes were obtained from BnIR (https://yanglab.hzau.edu.cn/). The phylogenetic trees were constructed using MEGA_v7.0.21 software (Kumar et al. 2016) using the neighbor-joining method and a bootstrap test that was replicated 1000 times.

### Statistics

Statistical significance was determined using R (https://r-project.org). The Wilcoxon rank sum test and the χ2 test were performed using the *Wilcoxon. test* function and the *chisq. test* function, respectively, from the R package.

## Ethical statement

This article does not contain any studies with human or animal subjects.

## Declaration of competing interest

The authors declare that they have no known competing financial interests or personal relationships that could have appeared to influence the work reported in this paper.

## Acknowledgments

We thank the National Key Laboratory of Crop Genetic Improvement of Huazhong Agricultural University for providing the bioinformatics computing platform, and Novogene provides sequencing services. This work was supported by the Science and Technology Innovation 2030-Major Project (2023ZD04068) to Cheng Dai and the National Natural Science Foundation of China (No. 32172070) to Chaozhi Ma.

## Author contributions

**D.C., Q.C., M.C., and D.S.** conceived the original idea. **D.S., W.L.,** and **L.R.** performed experiments; **Q.C.** and **D.S.** designed the work and analyzed the data; **Q.C.** and **D.C.** wrote the paper and discussed it with all authors.

## Data availability

All high-throughput sequencing data for RNA-seq, small RNA-seq, ATAC-seq, and WGBS of *Brassica* allotriploid hybrids and parents in this study are available in the Short Read Archive (SRA) under NCBI BioProject accession number PRJNA1154317. All high-throughput sequencing data for sRNA-seq and WGBS of *Bnadcl3*^CR^ and *Bnardr2*^CR^ mutants are available in SRA, NCBI BioProject accession number PRJNA1157971.

## SUPPLEMENTAL INFORMATION

### Supplemental Figure legends

**Figure S1. Schematic model for the diploid species (*B. rapa*, Hort), F_1_ hybrids, and the allotetraploid species (*B. napus*, *s70*, and *yu25*).** The images showed the phenotypes of the allotriploid F_1_ hybrids and their parental lines at the flowering stage and the typical chromosome number (Scale bars = 2 µm) of the allotriploid *Brassica* hybrids (F_1_ hybrids).

**Figure S2. The bar graph showed the content of total flavonoids in maternal lines (*s70* and *yu25*), paternal line (Hort), and F_1_ hybrids (sh and yh).**

**Figure S3. Gene expression of the three subgenomes in F_1_ hybrids. (a)** Histograms showing the expression levels (fragments per kilobase per million, FPKM) of all genes, subgenomic unique genes, and homologous genes in the A_n_, C_n_, and A_r_ subgenomes in the 40-days (40D) and 60-days (60D) F_1_ hybrids. **(b)** Number of dominant homologous genes and suppressed homologous genes in the A_n_, C_n_, and A_r_ subgenomes of F_1_ hybrids and *in silico* ‘hybrids’.

**Figure S4. Distribution of ACRs among the three subgenomes. (a)** Chromatin accessibility around A_n_, C_n_ and A_r_ subgenome genes in F_1_ hybrids (Hybrid-sh and Hybrid-yh) and *in silico* ‘hybrids’ (*in silico*-sh and *in silico*-yh). **(b)** Percentage of dominant homologous triplet genes in the A_n_, C_n_ and A_r_ subgenomes in two hybrids. **(c)** These genes were divided into four patterns based on the ACR number of the homologous triplet genes:A_r_ = A_n_ ≠ C_n_, A_r_ = C_n_ ≠ A_n_, A_n_ = C_n_ ≠ A_r_, and A_r_ = A_n_ = C_n._ Histogram showing the number of each pattern.

**Figure S5. TEs methylation landscape of different subgenomes in F_1_ hybrids. (a-c)** DNA methylation levels of TEs **(a)**, short TEs **(b)**, and long TEs **(c)** in the A_n_, A_r_, and C_n_ subgenomes.

**Figure S6. DNA methylation profiles around the dominant and suppressed homologous triplet genes. (a-b)** DNA methylation profiles around dominant genes **(a)** and suppressed genes **(b)** in A_n_, C_n_ and A_r_ subgenomes.

**Figure S7. Genome-wide DNA methylation profile and expression of methyltransferase-related genes. (a)** Genome-wide DNA methylation profiles of A_n_, C_n_ and A_r_ subgenomes in Hybrid-yh and *in silico*-yh. **(b)** Phylogenetic analysis of DNA methyltransferase genes in *B. napus* and *B. rapa*. Heatmap showing DNA methyltransferase gene transcript levels in the Hybrid-yh and *in silico*-yh. *S* and *H* represent *in silico*-yh and Hybrid-yh, respectively. **(c)** Number of 24-nt siRNA clusters of the A_n_, C_n_ and A_r_ subgenomes in the Hybrid-yh and *in silico*-yh.

**Figure S8. DNA methylation levels in WT, *Bnadcl3^CR^*, and *Bnardr2^CR^* mutants. (a)** Profiles of methylation levels around WT and *Bnardr2^CR^* mutant genes in the A_n_ and C_n_ subgenomes. **(b)** Profiles of methylation levels around WT, *Bnadcl3^CR^* and *Bnardr2^CR^* mutant TEs. **(c)** Boxplots showing short TE (left) and long TE (right) methylation levels in WT, *Bnadcl3^CR^* and *Bnardr2^CR^* mutants. Average DNA methylation levels were determined from two biological replicates per genotype.

**Figure S9. Distribution of 24-nt siRNA clusters in *Bnadcl3^CR^* and *Bnardr2^CR^* mutants. (a)** Number of overlaps of *Bnadcl3^CR^* (left) and *Bnardr2^CR^* (right) mutant 24-nt siRNA clusters with WT. **(b)** Proportion of differentially expressed 24-nt siRNA clusters in long TEs and short TEs in *Bnadcl3^CR^* and *Bnardr2^CR^* mutants compared with WT. **(c)** Gene expression levels of demethylases. *Reactive Oxygen Species 1 (ROS1)*; *DEMETER (DME); DEMETER-LIKE 3 (ML3)*; *Increased DNA Methylation 1 (IDM1)*.

### Supplemental Tables

**Table S1.** Statistics of RNA-seq data and reads mapping for all samples.

**Table S2.** Statistics of ATAC-seq data and reads mapping for all samples.

**Table S3.** Statistics of WGBS data and reads mapping for all samples.

**Table S4.** Statistics of sRNA-seq data and reads mapping for all samples.

**Table S5.** The number of ACRs for all samples.

**Table S6.** Statistics of WGBS data and reads mapping for wild type and mutants.

**Table S7.** Statistics of sRNA-seq data and reads mapping for wild type and mutants.

**Table S8.** Genotype of the *Bnadcl3*^CR^ and *Bnardr2*^CR^ double mutants.

**Table S9.** The BnaRDR2 and BnaDCL3 target-specific primers.

## Notes

### Competing Interest Statement

The authors have declared no competing interest.

