## Supplemental table for "The dynamics of ACR and DNA methylation impact asymmetric subgenome dominance in allotriploid *Brassica* species"

**Title Page**

Chengtao Quan^1,3^, Tian Xia^1,3^, Lulin Wang^3,4^, Ruifan Liu^1,3^, Chaozhi Ma^1,3^, Shengwei Dou^2,*^, and Cheng Dai^1,3,*^

1 National Key Laboratory of Crop Genetic Improvement, Huazhong Agricultural University, Wuhan 430070, China

2 College of Horticulture Science and Engineering, Shandong Agricultural University, Tai’an 271018, China

3 Hubei Hongshan Laboratory, Wuhan, 430070, China

4 National Key Laboratory for Germplasm Innovation and Utilization of Horticultural Crops, College of Horticulture and Forestry Sciences, Huazhong Agricultural University, Wuhan 430070, China

The author responsible for the distribution of materials integral to the findings presented in this article following the policy described in the Instructions for Authors are:

Shengwei Dou

Cheng Dai

To whom correspondence should be addressed.

Dr. Shengwei Dou

College of Horticulture Science and Engineering, Shandong Agricultural University, Tai’an 271018, P.R. China

Dr. Cheng Dai

National Key Laboratory of Crop Genetic Improvement, Huazhong Agricultural University, Wuhan 430070, P.R. China

**Supporting Information**

**Supplemental Figure legends**

**Figure S1.** Schematic model for the diploid species (*B. rapa*, Hort), F_1_ hybrids, and the allotetraploid species (*B. napus*, *s70* and *yu25*).

**Figure S2.** Determination of primary metabolites.

**Figure S3.** Gene expression of the three subgenomes in F_1_ hybrids.

**Figure S4.** Distribution of ACRs among the three subgenomes.

**Figure S5.** TEs methylation landscape of different subgenomes in F_1_ hybrids.

**Figure S6.** DNA methylation profiles around the dominant and suppressed homologous triplet genes.

**Figure S7.** Genome-wide DNA methylation profile and expression of methyltransferase-related genes.

**Figure S8.** DNA methylation levels in WT, *Bnadcl3^CR^,* and *Bnardr2^CR^* mutants.

**Figure S9.** Distribution of 24-nt siRNA clusters in *Bnadcl3^CR^* and *Bnardr2^CR^* mutants.

**Supplemental Tables**

**Table S1.** Statistics of RNA-seq data and reads mapping for all samples.

**Table S2.** Statistics of ATAC-seq data and reads mapping for all samples.

**Table S3.** Statistics of WGBS data and reads mapping for all samples.

**Table S4.** Statistics of sRNA-seq data and reads mapping for all samples.

**Table S5.** The number of ACRs for all samples.

**Table S6.** Statistics of WGBS data and reads mapping for wild type and mutants.

**Table S7.** Statistics of sRNA-seq data and reads mapping for wild type and mutants.

**Table S8.** Genotype of the *Bnadcl3*^CR^ and *Bnardr2*^CR^ double mutants.

**Table S9.** The BnaRDR2 and BnaDCL3 target-specific primers.

**Supplemental figure legends**

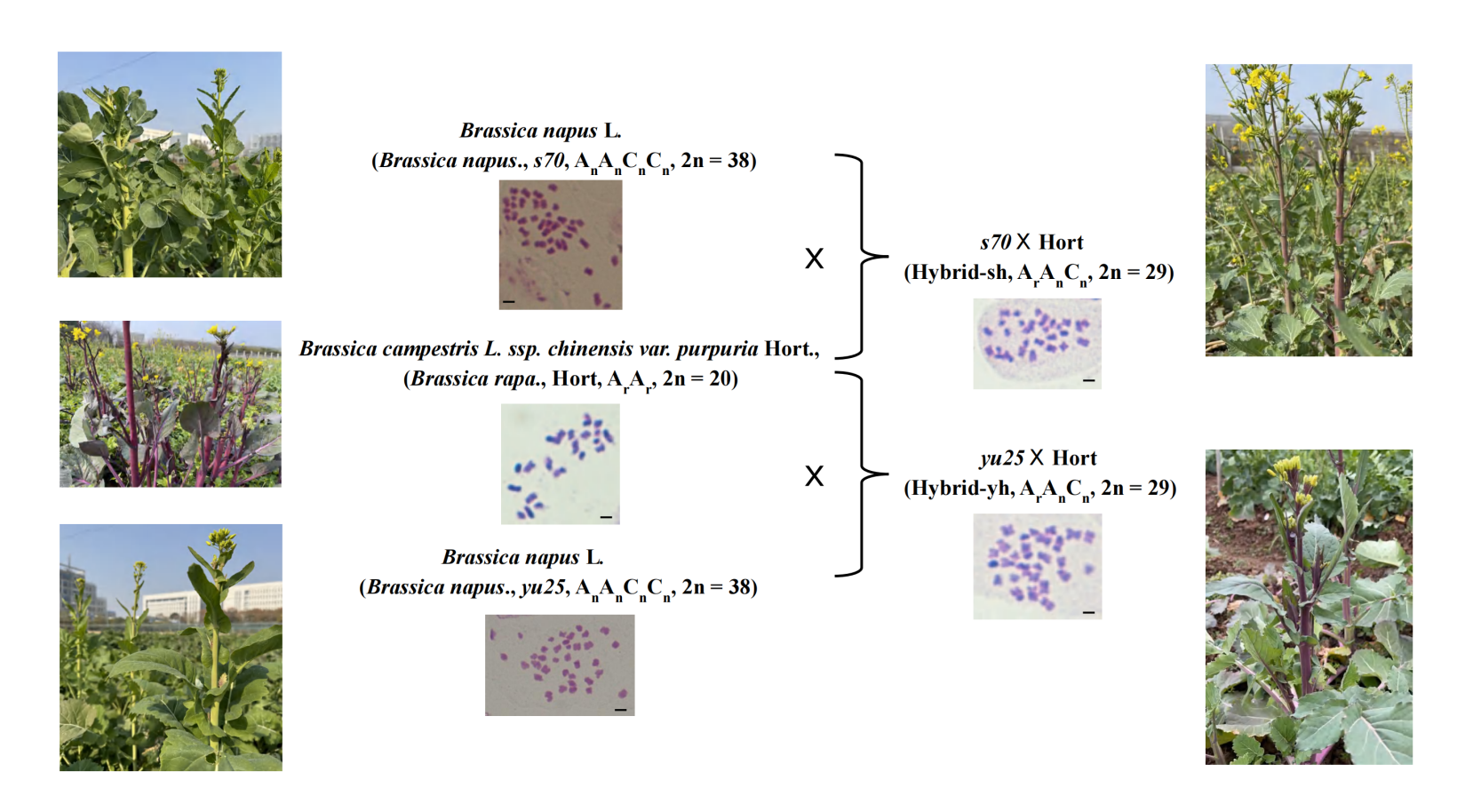

**Figure S1. Schematic model for the diploid species (*B. rapa*, Hort), F_1_ hybrids, and the allotetraploid species (*B. napus*, *s70* and *yu25*).** The images showed the phenotypes of the allotriploid F_1_ hybrids and their parental lines at the flowering stage and the typical chromosome number (Scale bars = 2 µm) of the allotriploid *Brassica* hybrids (F_1_ hybrids).

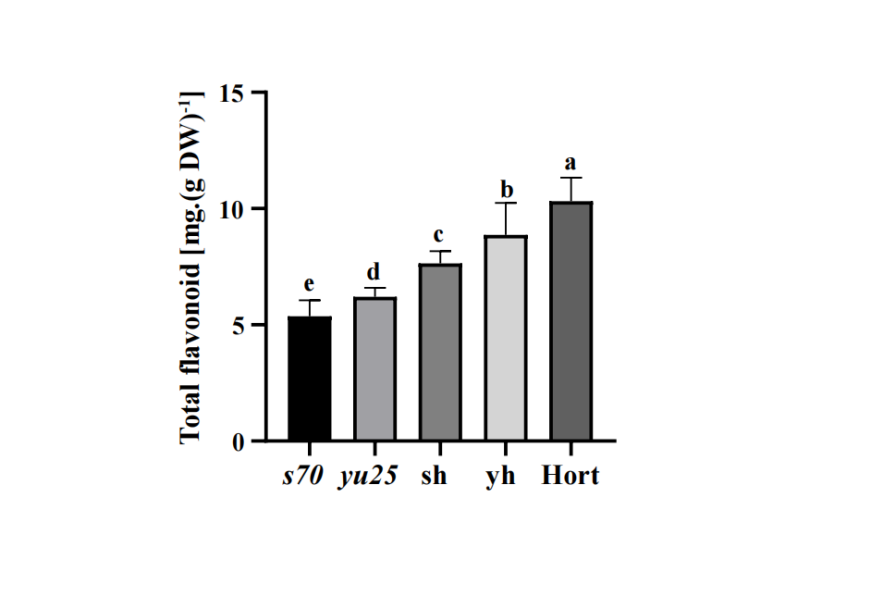

**Figure S2. The bar graph showed the content of total flavonoids in maternal lines (*s70* and *yu25*), paternal line (Hort), and F_1_ hybrids (sh and yh).**

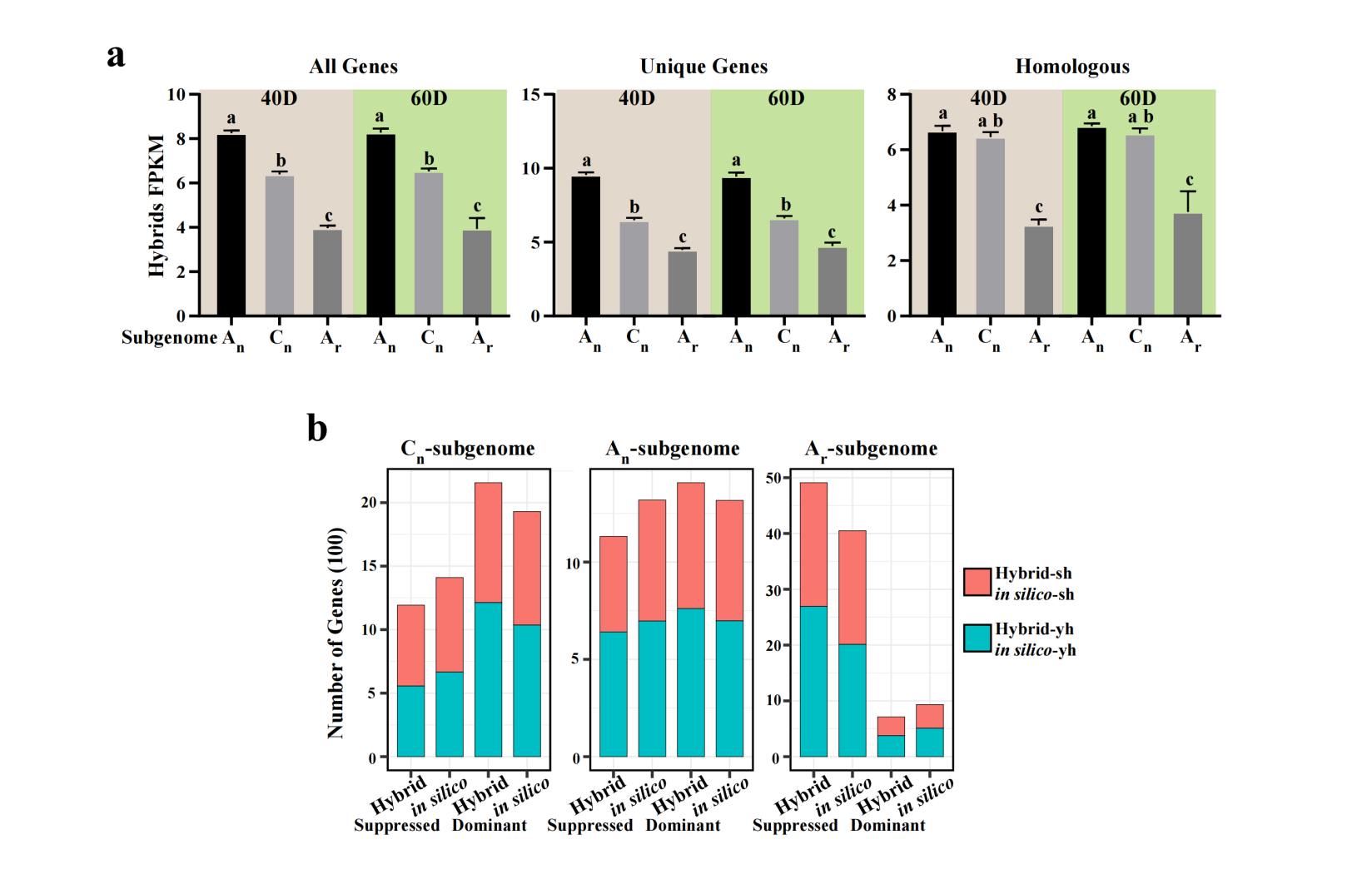

**Figure S3. Gene expression of the three subgenomes in F_1_ hybrids. (a)** Histograms showing the expression levels (fragments per kilobase per million, FPKM) of all genes, subgenomic unique genes, and homologous genes in the A_n_, C_n_, and A_r_ subgenomes in the 40-days (40D) and 60-days (60D) F_1_ hybrids. **(b)** Number of dominant homologous genes and suppressed homologous genes in the A_n_, C_n_, and A_r_ subgenomes of F_1_ hybrids and *in silico* 'hybrids'.

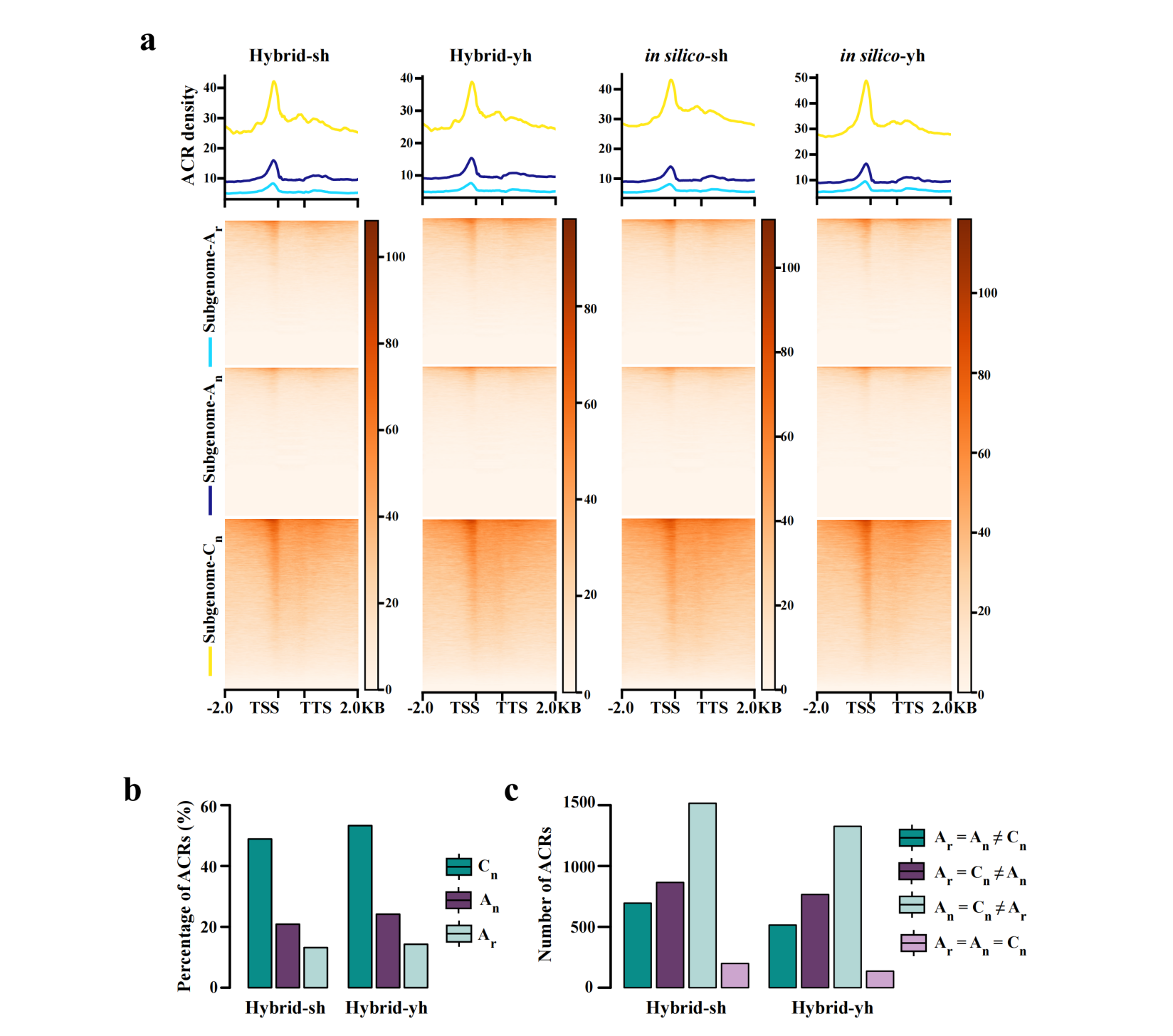

**Figure S4. Distribution of ACRs among the three subgenomes. (a)** Chromatin accessibility around A_n_, C_n_ and A_r_ subgenome genes in F_1_ hybrids (Hybrid-sh and Hybrid-yh) and *in silico* 'hybrids' (*in silico*-sh and *in silico*-yh). **(b)** Percentage of dominant homologous triplet genes in the A_n_, C_n_ and A_r_ subgenomes in two hybrids. **(c)** These genes were divided into four patterns based on the ACR number of the homologous triplet genes:A_r_ = A_n_ ≠ C_n_, A_r_ = C_n_ ≠ A_n_, A_n_ = C_n_ ≠ A_r_, and A_r_ = A_n_ = C_n._ Histogram showing the number of each pattern.

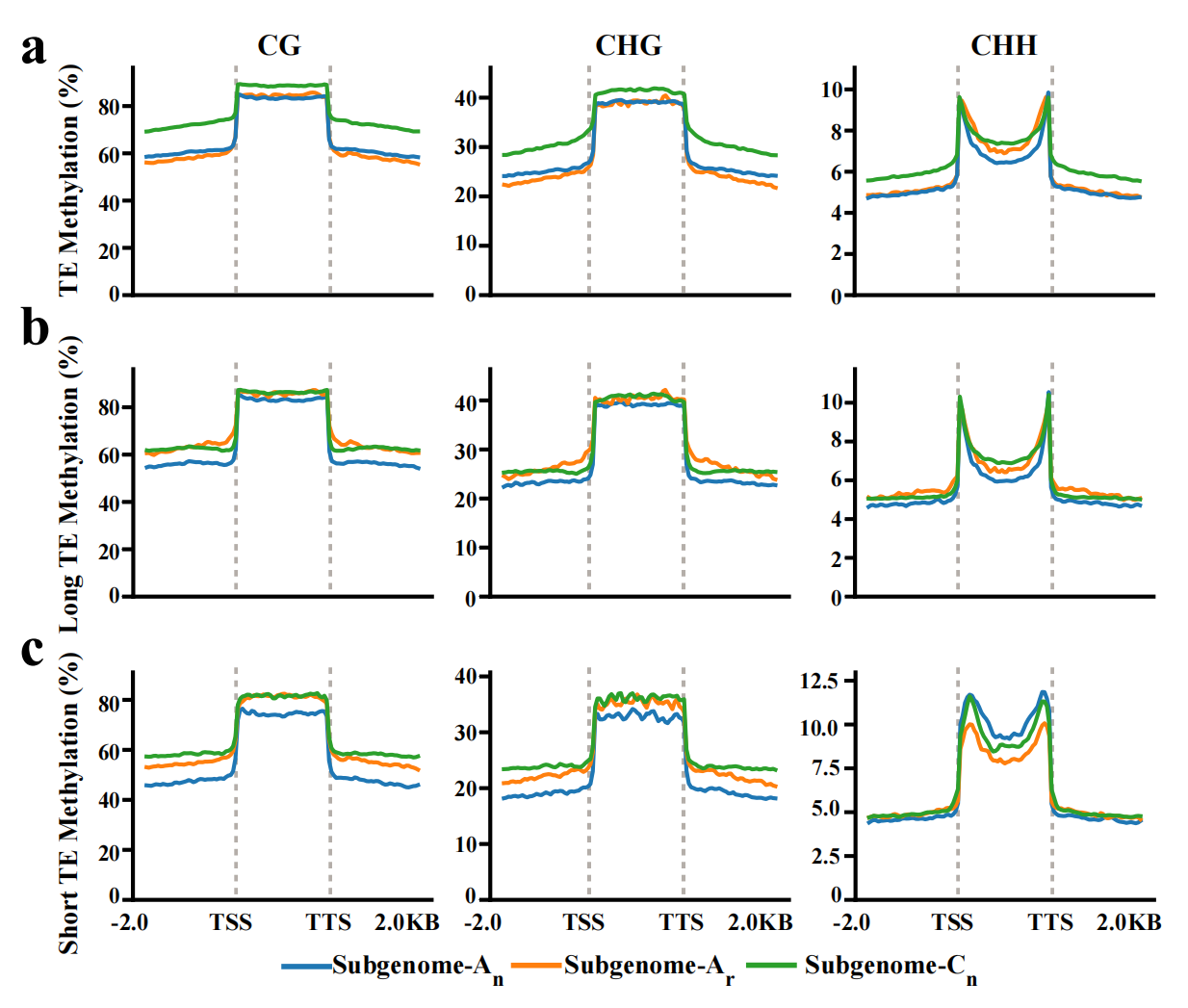

**Figure S5.** **TEs methylation landscape of different subgenomes in F_1_ hybrids.** **(a-c)** DNA methylation levels of TEs **(a)**, short TEs **(b)**, and long TEs **(c)** in the A_n_, A_r_, and C_n_ subgenomes.

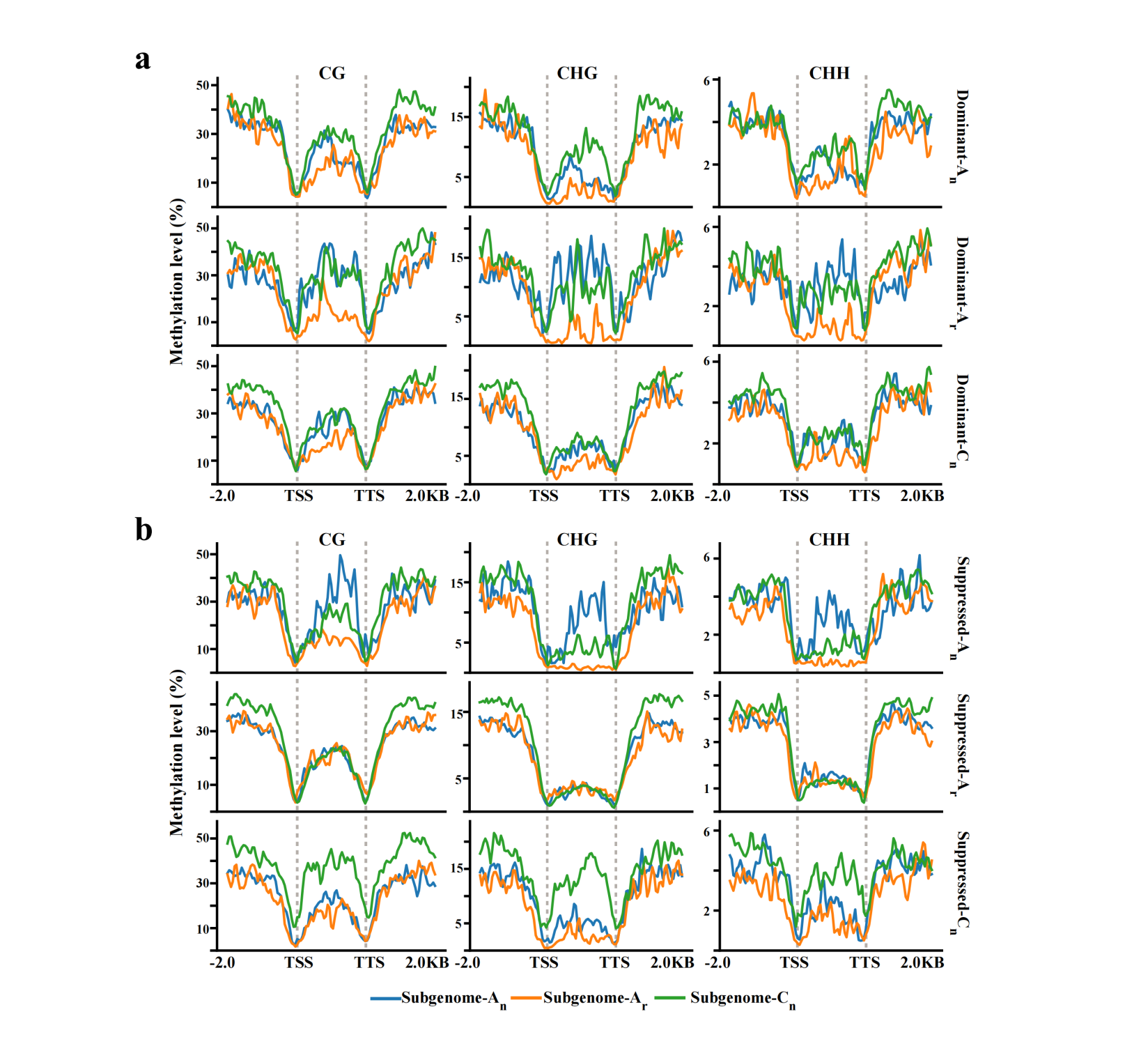

**Figure S6. DNA methylation profiles around the dominant and suppressed homologous triplet genes. (a-b)** DNA methylation profiles around dominant genes **(a)** and suppressed genes **(b)** in A_n_, C_n_ and A_r_ subgenomes.

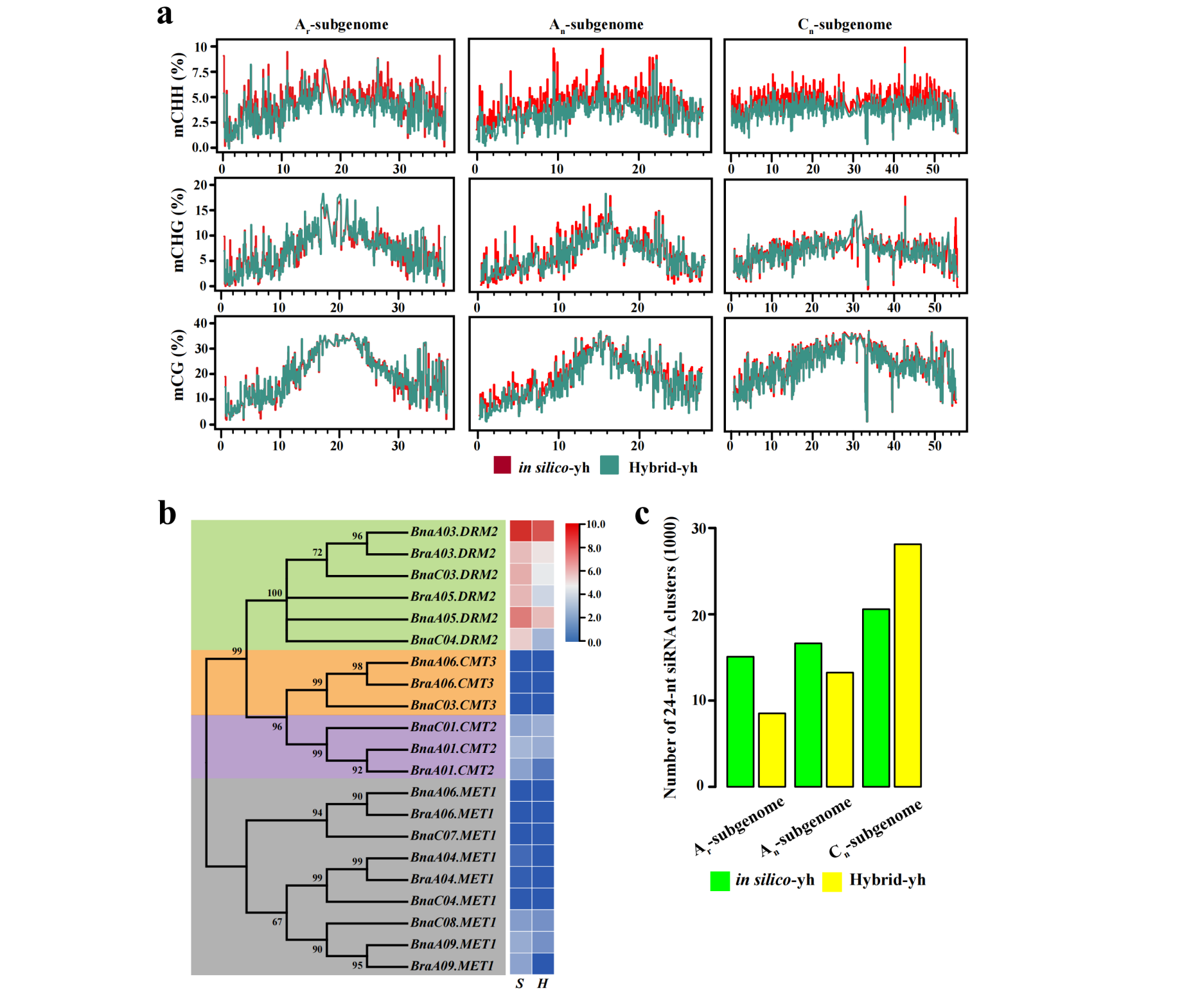

**Figure S7. Genome-wide DNA methylation profile and expression of methyltransferase-related genes. (a)** Genome-wide DNA methylation profiles of A_n_, C_n_ and A_r_ subgenomes in the Hybrid-yh and *in silico*-yh. **(b)** Phylogenetic analysis of DNA methyltransferase genes in *B. napus* and *B. rapa*. Heatmap showing DNA methyltransferase gene transcript levels in Hybrid-yh and *in silico*-yh. *S* and *H* represent *in silico*-yh and Hybrid-yh, respectively. **(c)** Number of 24-nt siRNA clusters of the A_n_, C_n_ and A_r_ subgenomes in the Hybrid-yh and *in silico*-yh.

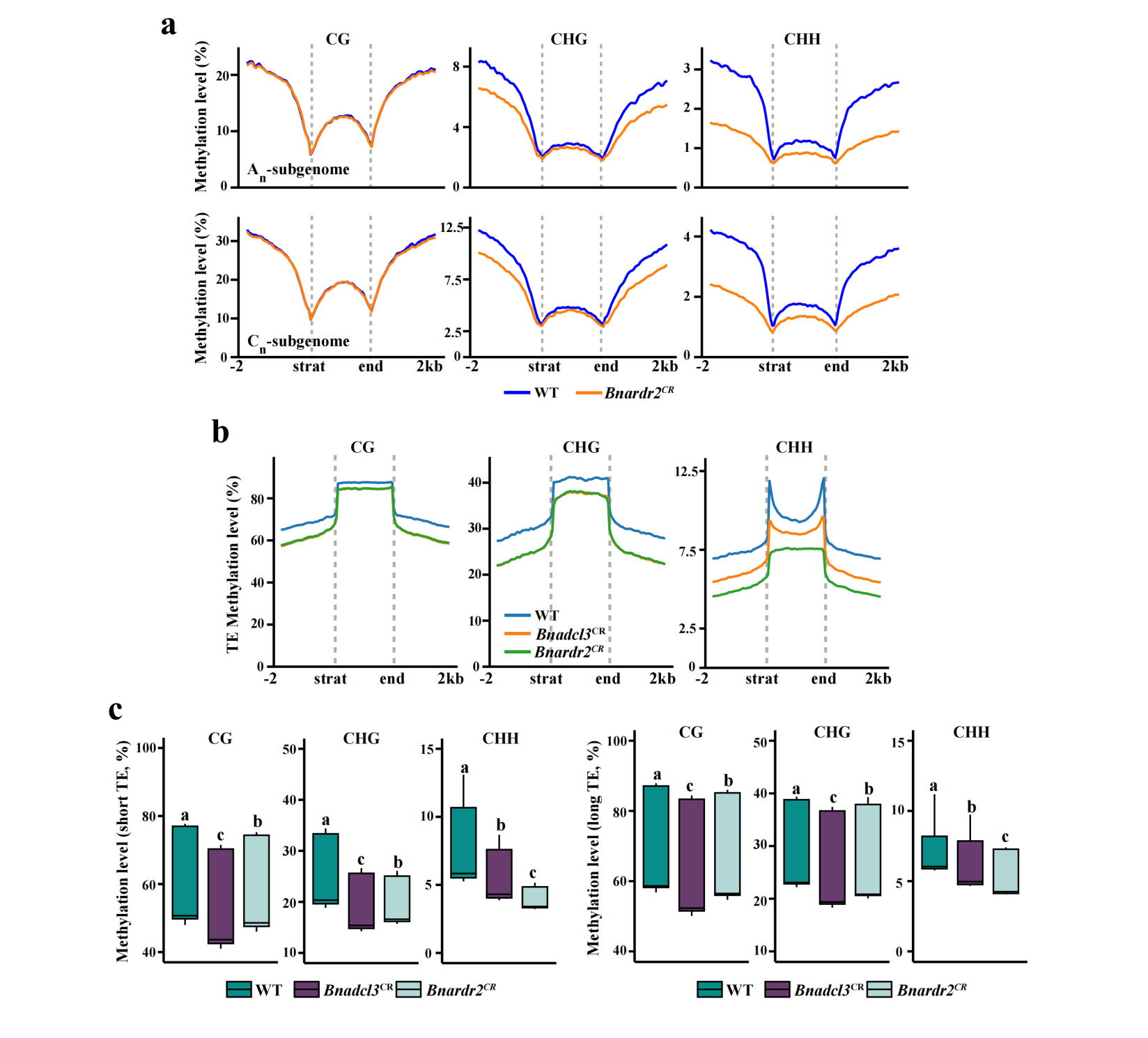

**Figure S8. DNA methylation levels in WT, *Bnadcl3^CR^*, and *Bnardr2^CR^* mutants. (a)** Profiles of methylation levels around WT and *Bnardr2^CR^* mutant genes in the A_n_ and C_n_ subgenomes. **(b)** Profiles of methylation levels around WT, *Bnadcl3^CR^* and *Bnardr2^CR^* mutant TEs. **(c)** Boxplots showing short TE (left) and long TE (right) methylation levels in WT, *Bnadcl3^CR^* and *Bnardr2^CR^* mutants. Average DNA methylation levels were determined from two biological replicates per genotype.

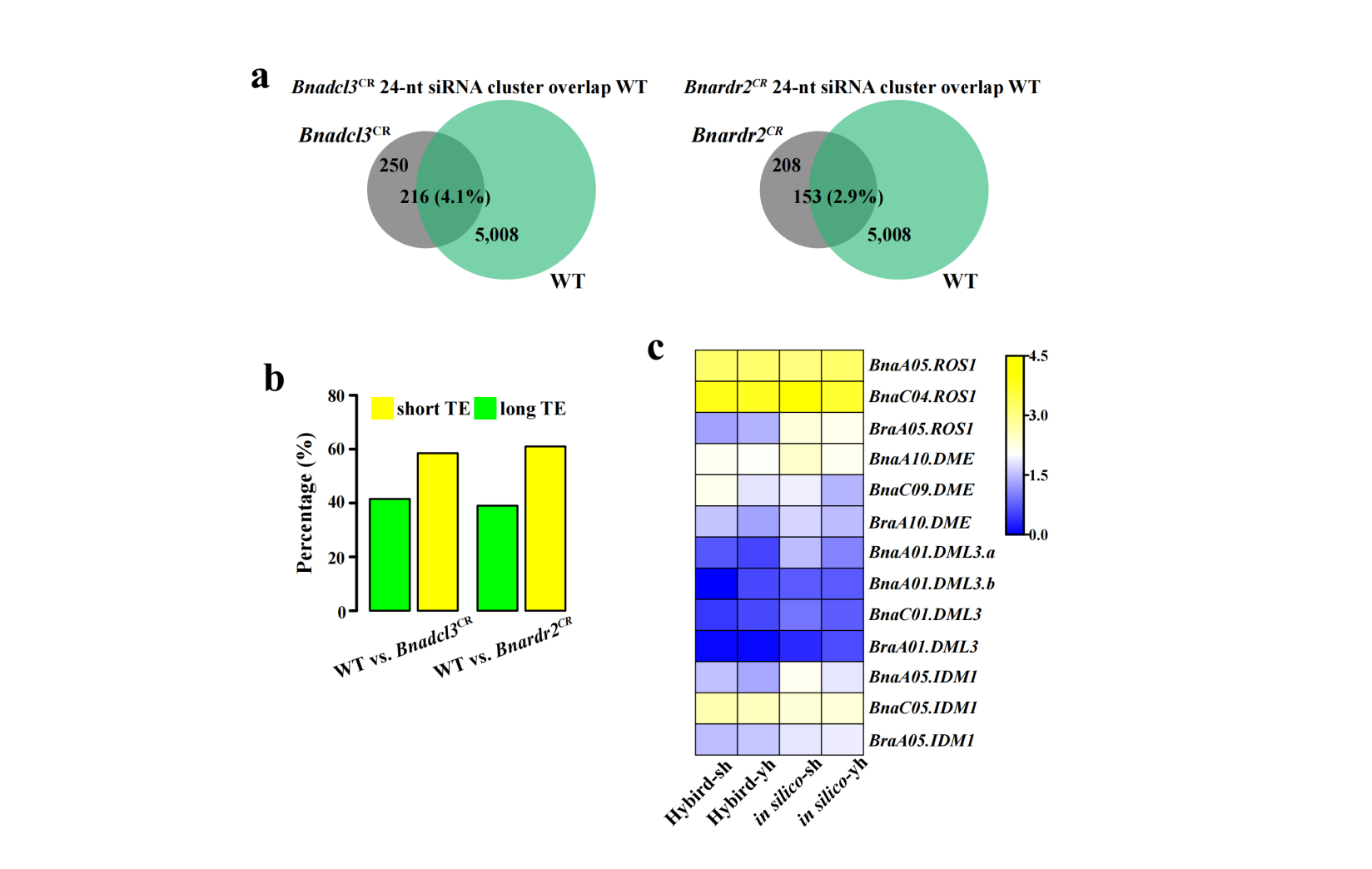

**Figure S9. Distribution of 24-nt siRNA clusters in *Bnadcl3^CR^* and *Bnardr2^CR^* mutants. (a)** Number of overlaps of *Bnadcl3^CR^* (left) and *Bnardr2^CR^* (right) mutant 24-nt siRNA clusters with WT. **(b)** Proportion of differentially expressed 24-nt siRNA clusters in long TEs and short TEs in *Bnadcl3^CR^* and *Bnardr2^CR^* mutants compared with WT. (**c**) Gene expression levels of demethylases. *Reactive Oxygen Species 1* (*ROS1*); *DEMETER* (*DME*); *DEMETER-LIKE 3* (*ML3*); *Increased DNA Methylation 1* (*IDM1*).

**Supplemental Table S1. Statistics of RNA-seq data and reads mapping for all samples.**

| **RNA-seq** | **Raw Reads**  **(millions)** | **Clean Reads**  **(millions)** | **Q30 (%)** | **GC Content (%)** | **Mapped Clean reads (millions)** | ***R*** |
| --- | --- | --- | --- | --- | --- | --- |
| ***s70*** | 20.67 | 19.21 | 93.65 | 47.31 | 18.02 | 0.96 |
| ***s70*** | 20.54 | 19.20 | 93.82 | 47.23 | 18.07 |  |
| ***s70*** | 21.95 | 20.82 | 93.57 | 46.45 | 19.57 |  |
| ***yu25*** | 21.72 | 20.54 | 93.22 | 47.51 | 19.37 | 0.95 |
| ***yu25*** | 21.37 | 20.13 | 93.05 | 47.2 | 18.99 |  |
| ***yu25*** | 21.01 | 20.54 | 93.39 | 46.7 | 19.55 |  |
| **Hybrid-sh** | 21.48 | 20.94 | 92.71 | 46.78 | 19.84 | 0.95 |
| **Hybrid-sh** | 21.43 | 20.58 | 93.19 | 46.38 | 19.64 |  |
| **Hybrid-sh** | 19.78 | 18.71 | 93.78 | 46.81 | 17.65 |  |
| **Hybrid-yh** | 19.76 | 18.59 | 93.19 | 46.86 | 17.64 | 0.97 |
| **Hybrid-yh** | 19.90 | 18.86 | 93.09 | 47.3 | 17.82 |  |
| **Hybrid-yh** | 21.25 | 20.00 | 93.38 | 47.19 | 18.98 |  |
| **Hort** | 21.86 | 20.82 | 93.21 | 47.27 | 18.50 | 0.96 |
| **Hort** | 22.98 | 21.88 | 93.65 | 47.3 | 19.46 |  |
| **Hort** | 21.64 | 20.42 | 93.59 | 47.35 | 18.10 |  |

**Supplemental Table S2. Statistics of ATAC-seq data and reads mapping for all samples.**

| **ATAC-seq** | **Raw Reads**  **(millions)** | **Clean Reads**  **(millions)** | **Q30(%)** | **SPOT** | **Mapped Clean reads (millions)** | ***R*** |
| --- | --- | --- | --- | --- | --- | --- |
| ***s70*** | 46.71 | 46.41 | 94.46 | 0.28 | 41.92 | 0.94 |
| ***s70*** | 48.94 | 48.63 | 92.02 | 0.26 | 37.99 |  |
| ***s70*** | 50.59 | 50.29 | 92.2 | 0.29 | 39.55 |  |
| ***yu25*** | 51.05 | 50.75 | 92.48 | 0.28 | 38.86 | 0.92 |
| ***yu25*** | 49.49 | 49.21 | 92.46 | 0.26 | 38.19 |  |
| ***yu25*** | 48.24 | 47.95 | 92.01 | 0.28 | 37.65 |  |
| **Hybrid-sh** | 44.18 | 43.85 | 91.57 | 0.28 | 31.21 | 0.95 |
| **Hybrid-sh** | 50.01 | 49.60 | 92.27 | 0.28 | 38.72 |  |
| **Hybrid-sh** | 50.51 | 50.12 | 92.34 | 0.26 | 38.39 |  |
| **Hybrid-yh** | 45.18 | 44.70 | 92.31 | 0.25 | 33.38 | 0.94 |
| **Hybrid-yh** | 48.94 | 48.48 | 91.77 | 0.28 | 38.49 |  |
| **Hybrid-yh** | 41.34 | 41.02 | 92.07 | 0.25 | 29.96 |  |
| **Hort** | 45.15 | 44.63 | 90.71 | 0.32 | 35.03 | 0.96 |
| **Hort** | 48.04 | 47.61 | 91.28 | 0.32 | 36.01 |  |
| **Hort** | 44.91 | 44.51 | 91.23 | 0.33 | 34.21 |  |

**Supplemental Table S3. Statistics of WGBS data and reads mapping for all samples.**

| **WGBS** | **QC-passed reads (millions)** | **Properly paired reads**  **(millions)** | **Average Cytosine depth** | **Bisulfite conversion (%)** | **mC/(C+T) (%)** | ***R*** |
| --- | --- | --- | --- | --- | --- | --- |
| ***s70*** | 203.75 | 192.64 | 30.31 | 99.58 | 10.61 | 0.91 |
| ***s70*** | 242.64 | 196.80 | 33.61 | 99.21 | 13.29 |  |
| ***yu25*** | 203.75 | 193.56 | 30.26 | 99.32 | 14.47 | 0.90 |
| ***yu25*** | 193.43 | 184.32 | 28.84 | 99.45 | 12.95 |  |
| **Hybrid-sh** | 199.42 | 188.15 | 29.66 | 99.26 | 12.52 | 0.89 |
| **Hybrid-sh** | 206.21 | 195.15 | 30.65 | 99.38 | 10.26 |  |
| **Hybrid-yh** | 184.64 | 173.47 | 27.41 | 99.41 | 12.03 | 0.92 |
| **Hybrid-yh** | 207.84 | 197.32 | 30.86 | 99.55 | 11.03 |  |
| **Hort** | 193.80 | 177.98 | 28.54 | 99.16 | 9.85 | 0.88 |
| **Hort** | 238.28 | 219.90 | 35.04 | 99.73 | 10.42 |  |

**Supplemental Table S4. Statistics of sRNA-seq data and reads mapping for all samples.**

| **sRNA-seq** | **Raw Reads**  **(millions)** | **Clean Reads**  **(millions)** | **Q20 (%)** | **Q30 (%)** | **Mapped Clean reads (millions)** | ***R*** |
| --- | --- | --- | --- | --- | --- | --- |
| ***s70*** | 29.80 | 28.64 | 99.3 | 97.7 | 26.04 | 0.95 |
| ***s70*** | 28.59 | 27.12 | 99.1 | 97.1 | 26.48 |  |
| ***yu25*** | 29.75 | 28.26 | 99.1 | 96.9 | 27.27 | 0.92 |
| ***yu25*** | 30.15 | 28.85 | 98.2 | 95.1 | 27.90 |  |
| **Hybrid-sh** | 29.15 | 27.03 | 98.4 | 95.6 | 26.16 | 0.92 |
| **Hybrid-sh** | 28.09 | 27.31 | 98.6 | 95.9 | 26.11 |  |
| **Hybrid-yh** | 28.39 | 27.74 | 98.5 | 95.7 | 26.82 | 0.93 |
| **Hybrid-yh** | 28.17 | 27.51 | 98.2 | 95 | 26.56 |  |
| **Hort** | 28.70 | 28.02 | 98.9 | 96.5 | 26.91 | 0.91 |
| **Hort** | 28.40 | 27.83 | 99.1 | 97.1 | 26.41 |  |

**Supplemental Table S5. The number of ACRs for all samples.**

| **Samples** | **Whole genome** | **Overlap** | **A_r_-genome** | **A_n_-genome** | **C_n_-genome** |
| --- | --- | --- | --- | --- | --- |
| ***s70*** | 45,661 |  |  | 16,698 | 28,963 |
| ***s70*** | 62,814 | 72,947 |  | 22,936 | 39,878 |
| ***s70*** | 84,194 |  |  | 31,681 | 52,513 |
| ***yu25*** | 49,084 |  |  | 17,955 | 31,129 |
| ***yu25*** | 51,107 | 61,316 |  | 18,457 | 32,650 |
| ***yu25*** | 55,622 |  |  | 19,737 | 35,885 |
| **Hybrid-sh** | 21,780 |  | 2,744 | 5,818 | 13,218 |
| **Hybrid-sh** | 37,125 | 31,046 | 4,814 | 9,536 | 22,775 |
| **Hybrid-sh** | 48,936 |  | 6,751 | 13,703 | 28,482 |
| **Hybrid-yh** | 42,409 |  | 5,533 | 10,869 | 26,007 |
| **Hybrid-yh** | 40,264 | 35,639 | 5,058 | 9,284 | 25,922 |
| **Hybrid-yh** | 33,413 |  | 4,244 | 8,182 | 20,987 |
| **Hort** | 31,357 |  | 31,357 |  |  |
| **Hort** | 30,543 | 38,553 | 30,543 |  |  |
| **Hort** | 29,862 |  | 29,862 |  |  |

**Supplemental Table S6. Statistics of WGBS data and reads mapping for wild type and mutants.**

| **WGBS** | **QC-passed reads (millions)** | **Properly paired reads**  **(millions)** | **Average Cytosine depth** | **Bisulfite conversion (%)** | **mC/(C+T) (%)** | ***R*** |
| --- | --- | --- | --- | --- | --- | --- |
| **WT** | 231.78 | 219.56 | 34.06 | 99.50 | 11.88 | 0.91 |
| **WT** | 190.56 | 182.47 | 28.99 | 99.23 | 14.79 |  |
| ***Bnadcl3*^CR^** | 243.95 | 230.61 | 35.85 | 99.73 | 11.29 | 0.92 |
| ***Bnadcl3*^CR^** | 208.89 | 197.01 | 30.71 | 99.56 | 11.03 |  |
| ***Bnardr2*^CR^** | 243.08 | 231.66 | 35.73 | 99.72 | 10.70 | 0.89 |
| ***Bnardr2*^CR^** | 216.31 | 205.91 | 31.77 | 99.56 | 12.49 |  |

**Supplemental Table S7. Statistics of sRNA-seq data and reads mapping for wild type and mutants.**

| **sRNA-seq** | **Raw Reads**  **(millions)** | **Clean Reads**  **(millions)** | **Q20 (%)** | **Q30 (%)** | **Mapped Clean reads (millions)** | ***R*** |
| --- | --- | --- | --- | --- | --- | --- |
| **WT** | 11.37 | 10.38 | 93.49 | 90.25 | 10.06 | 0.96 |
| **WT** | 12.18 | 11.03 | 94.26 | 91.54 | 10.61 |  |
| ***Bnadcl3*^CR^** | 12.92 | 11.69 | 94.24 | 91.34 | 11.21 | 0.93 |
| ***Bnadcl3*^CR^** | 12.84 | 11.11 | 94.24 | 91.46 | 10.44 |  |
| ***Bnardr2*^CR^** | 16.01 | 14.24 | 95.46 | 93.17 | 13.47 | 0.94 |
| ***Bnardr2*^CR^** | 16.48 | 12.61 | 95.88 | 94.00 | 12.14 |  |

**Supplemental Table S8. Genotype of the *Bnadcl3^CR^* and *Bnardr2^CR^* double mutants.**

| **T1** | | | |
| --- | --- | --- | --- |
|  |  | **sgRNA1** | **sgRNA2** |
| ***Bnadcl3^CR^-L5*** | **BnaA06.DCL3** | TGGTCATTGCGATTCACACCAGGAAGG (1i) | CCAGCC----TTGAAGTTGAAGCAGA (4d) |
|  | **BnaC03.DCL3** | TGGTCATTGCGATTCACACCTGGAAGG (1i) | CCAGCCTATTGTTGAAGTTGAAGCAGA (1i) |
| ***Bnardr2^CR^-L2*** | **BnaA09.RDR2** | **CCCCGTAAGATCCCAGGAGAAGGCTCG (1i)** | **AGG------------------------------CGT (30d)** |
|  | **BnaC09.RDR2** | **CCCCGT-----CCAGGAGAAGGCTCG (5d)** | **TTTTGTGCCCTAGAGAAATGTGGAAGG (1i)** |

Note: The protospacer adjacent motif (PAM), sgRNAs and mutations were marked by green, red, and blue, respectively. i: insertion; d: deletion.

**Supplemental Table S9. The BnaRDR2 and BnaDCL3 target-specific primers.**

|  | **Gene_id** | **Target sequence** |  |  |
| --- | --- | --- | --- | --- |
| ***BnaRDR2*** | **BnaA09g22040D** | **TGTGCCCTAGAGAAATGGGA AGG** | **BnRDR2-gR1-F** | **TA GGTCTCCTAGAGAAATGGGA**  **gttttagagctagaa** |
|  |  |  | **BnRDR2-gR1-R** | **AT GGTCTCATCTAGGGCACAt**  **gcaccagccgggaa** |
| ***BnaRDR2*** | **BnaCnng57100D** | **GCCTTCTCCTGGGATCTACG GGG** | **BnRDR2-gR1-F** | **TA GGTCTCCCTGGGATCTACG**  **gttttagagctagaa** |
|  |  |  | **BnRDR2-gR1-R** | **AT GGTCTCACCAGGAGAAGGC**  **tgcaccagccgggaa** |
| ***BnaDCL3*** | **BnaA06g19710D** | **GCTTCAACTTCAACAATGGC TGG** | **BnDCL3-gR1-F** | **TA GGTCTCCTTCAACAATGGC**  **gttttagagctagaa** |
|  |  |  | **BnDCL3-gR1-R** | **AT GGTCTCATGAAGTTGAAGC**  **tgcaccagccgggaa** |
| ***BnaDCL3*** | **BnaC03g54010D** | **TCATTGCGATTCACACCGGA AGG** | **BnDCL3-gR1-F** | **TA GGTCTCCATTCACACCGGA**  **gttttagagctagaa** |
|  |  |  | **BnDCL3-gR1-R** | **AT GGTCTCAGAATCGCAATGA**  **tgcaccagccgggaa** |
